# Body posture, gaze, and predator detection: how stalking behaviour may influence colour vision evolution

**DOI:** 10.1101/2023.05.30.542645

**Authors:** Pedro Z. de Moraes, Pedro Diniz, Daniel M. A. Pessoa

## Abstract

The success of a predatory attack is related to how much a predator manages to approach a prey without being detected. Some carnivore mammals use environmental objects as visual obstacles during stalking behaviour, allowing them to get closer to their prey while only showing parts of their coat or faces during movement or visual monitoring. Here, we investigate the influence of carnivores’ body postures and gaze on their detection by potential prey. To do so, we photographed taxidermized carnivore models (cougar, ocelot and lesser grison) in natural scenes and presented them to human dichromats (i.e., colourblind) and trichromats (i.e., normal colour vision). Our findings highlight the importance of predators’ complete body outline and gaze as search images during predator detection tasks. We also demonstrate how the coat and facial colour pattern of predators may camouflage their body outline and gaze, hampering predator detection. Furthermore, we observed that carnivore coat colour patterns may serve as an additional cue for trichromats, particularly in hidden carnivore detection tasks that proved to be more difficult for dichromats. We discuss our results within the context of a predator-prey arms race scenario, considering the evolutionary processes that may have generated the evidence presented in this study.

## Introduction

The ability to accurately infer environmental predation risk is fundamental for the survival of animals, given the drastic adaptive consequences of not responding adequately to a predatory attack [1–3]. Predators commonly use cryptic behaviours while trying to capture a prey (i.e., stalking behaviour), moving stealthily and using visual obstacles (e.g., vegetation) to hide their bodies [4–7], in order to get as close as possible to the prey and increase the chances for a successful attack [8]. Therefore, natural selection should favour prey that visually detect predators at the farthest possible distance, that adequately assess predation risk after detection, and that respond accordingly to risk level, ultimately avoiding the predatory attack [1-3,9].

The first obstacle to the visual detection of a predator is the colour pattern presented by its surface. Several types of colour camouflage strategies evolved in animals [10], such as background-matching, in which predators’ colours resemble the colours of the background environment, or disruptive colouration, in which predators’ colour patterns create internal false edges that hamper the visual identification of their body in the environment [11–13]. These camouflage strategies interfere on prey visual processing by adding irrelevant or false information (i.e., noise) through the use of colour patterns and false edges, which in turn make it difficult to visually identify the predator (i.e., the signal) by minimizing signal-to-noise ratio [14]. In addition to colour, the predator’s body posture is especially important since it can provide additional information about actual predation risk [5,15,16]. Natural selection should favour animals that readily detect predators moving towards them, while selection for readily responding to predator moving away should be comparatively weaker, given that the former situation offers greater predation risk than the latter [5,17].

An even more informative visual cue to imminent predation risk is the direction of the predator’s gaze. Upon detecting a pair of eyes in the environment, a prey can assume that it is being observed by a predator and that it is in a context of high predation risk [18]. Several studies have demonstrated that vertebrates can detect gaze direction of potential predators [19–24] and respond accordingly. Gaze detection is especially relevant for primates, which also use this ability outside of predatory contexts, such as during intraspecific communication [18,25,26]. A possible evolutionary response to counteract gaze detection by prey is the selection of facial colouration patterns that either break predator’s facial features or camouflage predators’ eyes [5,27,28].

Given the interaction between colour patterns and visual cues indicative of predation risk (i.e., body posture and gaze), it is important to investigate how these factors may affect predator detection. Stalking behaviours are normally performed by mammals of order Carnivora, especially by members of family Felidae, and these commonly prey on other mammals [29–31]. The standard mammalian colour vision system is dichromacy, in which retina presents two different types of cones, each with a photopigment that maximally absorbs light at different wavelengths [32]. This visual system gives its bearer a colour vision similar to colourblind humans, with a limited perception of colours in bluish and yellowish hues. It is assumed that coat colouration patterns of carnivorous mammals were selected for camouflaging these animals against the colour vision system of their most common prey (i.e., other mammals), which are dichromats [33].

An exception to the mammalian visual pattern is found among primates. In this order, trichromacy (i.e., three different types of cones in the retina [34]) evolved in two distinct genetic contexts [35]. Primates of the parvorder Catarrhini (i.e., African and Asian primates) have a routine trichromacy, in which all individuals generally show the trichromatic visual system [36]. The Catarrhini have an autosomal gene which encodes a photopigment that absorbs short-wavelength light and two distinct genes, each encoding photopigments that absorb medium or long wavelengths [32,34]. Meanwhile, the parvorder Platyrrhini (i.e., primates from the Americas) typically shows a visual polymorphism in which members of the same species can be dichromats or trichromats [37]. In this group, the gene encoding medium/long absorption photopigments located on the X chromosome is polyallelic, so that heterozygous females are trichromats, while homozygous females and all males are dichromats [32,34].

Several hypotheses try to explain the evolution of a routine trichromacy in Catarrhini and a visual polymorphism in Platyrrhini. These hypotheses address relative advantages of each visual phenotype during the detection of food items (e.g., fruits, young leaves, flowers, insects) [38–55] or during intraspecific communication [56–60]. Despite the high number of research investigating the environmental factors that may have influenced primate colour vision evolution, only three studies experimentally evaluated the importance of predation risk on this matter to date [33,61,62]. Several studies have shown that the effectiveness of a camouflage strategy depends on the visual system that observes it [14,33,63–66]. As predators use colouration patterns to hide parts of their body, animals with more restricted colour vision might ignore chromatic components (e.g., the predator’s pelage colour) and focus on achromatic components (e.g., the outline of the predator’s body). Thus, prey with more limited colour vision (i.e., dichromats) would have an advantage in detecting camouflaged targets [63,65,66] compared to prey with more accurate colour vision (i.e., trichromats). However, Pessoa *et al.* [61] and de Moraes *et al.* [62] demonstrated through behavioural experiments that trichromats actually have an advantage in predator detection compared to dichromats. This advantage is even more accentuated in contexts of increased detection difficulty [62], such as predator detection in backgrounds of high visual complexity (e.g., forests), detection of predators with cryptic pelage colours, and long-distance predator detection.

The aim of the present study is to deepen the findings of Pessoa *et al.* [61] and de Moraes *et al.* [62] and investigate further visual cues that may be relevant to prey while they are scanning for predators. Specifically, we intend to evaluate how body posture and gaze of predators interact with the colour of their pelages and influence their detection by trichromats and dichromats. As previously demonstrated by de Moraes *et al.* [62], background environment and detection distance influence predator detection and, therefore, will also be considered in this study. First, we predict that the performance of both visual phenotypes should be similar on easier tasks, when obvious cues to the predator’s presence are available (e.g., full body outline or pair of eyes facing the observer). Second, we expect trichromats to be faster and more accurate, compared to dichromats, in more difficult tasks, that is, when predators present conspicuous pelage (i.e., yellow or orange colours) and their bodies are partially covered by background vegetation, or carnivores are turning their backs on the observer (i.e., not showing their face and eyes). Third, we predict that the performance of trichromats and dichromats will also be similar in extremely difficult tasks, such as detecting cryptically patterned carnivores, which have their bodies covered by vegetation and are positioned farthest away from the observer.

To test whether facial colouration patterns are able to camouflage the eyes and mask the gaze of predators [5,27], we expect that the performance of dichromats and trichromats will be similar when detecting forward-facing predators with uniformly coloured faces (i.e., whole face presents only one colour). In contrast, trichromacy will be advantageous when detecting predators that present complex facial colour patterns (i.e., face presents rosettes, lines, or patches of different colours). Finally, regardless of the observers’ visual phenotype, we expect predators displaying their full lateral body outline to be more easily detected than predators facing the observer and clearly showing their gaze (i.e., frontally positioned). This prediction is based on the fact that the lateral contour of the predator’s body may activate a larger area of the observer’s retina [67,68], resulting in a faster and more accurate response, compared to the smaller area activated by the predator’s frontal plane.

## Methods

### Experimental subjects

We used Ishihara (24 Plate Edition) and Hardy, Rand and Rittler (HRR 4th Edition, Laminated Version) pseudoisochromatic tests to select human dichromats and trichromats as experimental subjects (for more details on this protocol c.f., Moraes *et al.* [62]). All tests were applied during the day under indirect sunlight. All selected trichromats presented “normal readings” for all Ishihara plates, while all selected dichromats presented 16 “abnormal readings” on both the Transformation and Vanishing plates of the same test. We used the HRR test to confirm Ishihara’s diagnosis, and ultimately identified a total of 20 trichromats and 19 dichromats who agreed to participate in the experimental procedure (described below). All subjects were male undergraduate or graduate students between 18 and 30 years old. They all had normal visual acuity (20/20 vision), or corrected vision through the use of glasses or contact lenses when necessary. All dichromat individuals knew they were colourblind before we applied the pseudoisochromatic tests to them.

The experimental protocols for this study were approved by the Research Ethics Committee of Universidade Federal do Rio Grande do Norte (CEP/UFRN: number 026/11) and consent forms were signed by all participants. The documents referring to the identity of the subjects and their visual condition are archived at the Laboratory of Sensory Ecology of Universidade Federal do Rio Grande do Norte and only the authors have access to these files.

### Carnivore models

The taxidermized mammals used as stimuli in the detection tasks of this study were loaned to us by the Department of Zoology of Universidade de Brasilia and by the Museu de Ciencias Morfologicas of Universidade Federal do Rio Grande do Norte. The specimens were artistically taxidermized in neutral posture and their pelages were in good condition. A total of five taxidermized models were used, all of order Carnivora: one model of a cougar (*Puma concolor*), three ocelots (*Leopardus* spp.), and one lesser grison (*Galictis cuja*).

These models were chosen because they present three different types of pelage colour patterns. The cougar has a uniform reddish colouration that may have been selected as a camouflage strategy in open environments with high luminosity [69]. Ocelots have a dark-spotted yellowish pelage that may difficult the detection of these animals in habitats with dense vegetation and low luminosity [69]. The lesser grison have black abdomen and snout, while the top of its head and its dorsum have a light colouration reminiscent of dry grass, a colouration pattern known as light cape [69]. Lesser grisons also have a white line just above the eyes that separates the light (top) and dark (bottom) colour patches on their faces, which may be a type of disruptive colouration that creates false internal edges and hinders the whole visualization of animal’s head [12].

Although only ocelots are recognized primate predators [70–72], cougars and lesser grisons are predators of mammals [73,74] and therefore their pelages may have been selected as camouflage against dichromatic observers [33]. Thus, the carnivore models used in the experiments represent three levels of visual detection difficulty: (1) the cougar as the easiest visual detection model, with its conspicuous uniform reddish pelage; (2) the ocelots as an intermediate difficulty detection level, with their dark-spotted yellowish pelages; and (3) the lesser grison as hardest visual detection model, with its cryptic pelage [62].

The facial colour pattern presented by carnivore models also configure three difficulty levels regarding gaze detection. Cougars have a uniform facial colouring, so their eyes are easily identified on their face. Ocelots have dark spots and lines on their faces, highly contrasting with the yellowish background, which may difficult the recognition of facial features such as gaze. Lesser grisons present two colour patches on their face (black on the bottom and grey on the top), separated by a white line above the eyes. This lesser grison’s facial colouration pattern may hinder gaze detection since the dark eyes are surrounded by dark fur, forming a black patch. This, in turn, strongly contrasts with the white line above the eyes, which gives the impression that their face is not a uniform and easily recognizable structure but instead made up of two distinct objects. Therefore, we expect carnivore models’ faces to exert three difficulty levels in potential predator detection tasks: (1) cougar, as the model of easiest gaze detection because it does not present any complex colour pattern on its face [28]; (2) ocelots, as the models of intermediate gaze detection difficulty due to the black patches on their face that may hinder eye identification [5]; and (3) lesser grison, as the model of hardest gaze detection due to the disruptive colouration present on its face [12].

### Experimental stimuli

We positioned the carnivore models in backgrounds of natural vegetation and photographed these scenes to serve as experimental stimuli in the potential predator detection tasks. The photographs were taken in two different conserved Brazilian areas: Parque das Dunas, in the state of Rio Grande do Norte [05°48’S, 35°11’W] and Estacao Experimental Fazenda Agua Limpa of Universidade de Brasilia, in the Federal District [15°31’S, 47°42’W]. We received authorization from entities responsible for each area to use their grounds to take the photographs. We captured the photos using three different background scenarios: (1) grassland, an area of high luminosity where grasses predominate, and few trees and shrubs are present; (2) savanna, an area of high luminosity where the predominant vegetation is composed by shrubs and medium-sized trees; and (3) forest, an area of low luminosity, with random incidence of light and shade, and where tall trees and lianas predominate. Given the lighting and complexity of these environments, the background scenarios captured in the photographs add three levels of difficulty to the detection tasks: (1) grassland is the easiest detection scenario, with high luminosity and low visual complexity; (2) savanna is the intermediate difficulty detection scenario, with high luminosity and high visual complexity; and (3) forest is the hardest detection scenario, presenting low luminosity and high visual complexity [62].

We used an SRL digital camera (T2i 18 megapixel, Canon Inc.) to capture the photographs, and a tripod (Deluxe Tripod 200, Canon Inc.) to ensure camera stability. We took the pictures following the sequential method of Bergman & Beehner [75], in which photographs containing a colour chart (Pantone ColorChecker Passport, X-Rite Inc.) are taken after capturing an image that will serve as a positive (i.e., photo containing a carnivore model amid background vegetation) or negative (i.e., photo without a carnivore model, showing only background vegetation) stimuli in the experiment. It is important to keep an interval of one minute or less between the two photographs to ensure that stimuli photos and colour chart photos are taken under the same lighting conditions [75]. We used colour chart photos only for correction and standardization of colours present in stimuli photographs.

We followed the camera settings of previous studies that also used digital photographs to study animal colouration [61,62,76]. In summary, we set camera to AV mode (automatic shutter speed) with an ISO of 100 or 200, depending on available light. We used EF-S 18-55 mm lens (Canon, Inc) and set them at 55 mm focal length (88 mm equivalent). We set aperture to f/36 to obtain the maximum field depth. The decrease in sharpness caused by small aperture was not considered critical. We captured photographs in sunny cloudless days, between 9:00 and 15:30. We positioned the camera with our backs to the sun, so the camera always faced the sunlit side of carnivore models.

We created four body posture categories to simulate different types of movement of a potential predator (Fig. 1). In the first category, we positioned the carnivore model in the background environment with one side of its body facing the camera, so that the animal’s full lateral outline was visible in the photograph. In this posture, the model’s head faced the left or the right side of the image, and never faced the camera (i.e., both eyes were never visible to the observer). In the second category, we positioned the carnivore model facing the camera so that its face and eyes were fully visible to the observer. In this posture, only the model’s head, shoulders, and front legs were visible in the photo; we used scenery objects, such as bushes, logs and lianas, to hide the posterior portion of the body when necessary. In the third category, we positioned the carnivore model with its back to the camera, so that only its dorsum, hind legs, and tail were visible in the photo. We also used scenery objects to obstruct the anterior portion of the body when necessary. Finally, in the fourth category, we positioned the carnivore model with one of its sides facing the camera, but we visually obstructed parts of its body with objects from the scenery. Thus, the complete lateral outline of the animal’s body was unavailable to the observer, with only parts of its pelage being visible in photographs. With the four categories of body postures, we have added a new level of difficulty to potential predator detection tasks: (1) the lateral posture, presenting complete body outline, represents the task of easier detection; (2) the anterior posture, presenting face and eyes but not body complete outline, represents the second easiest task; (3) the posterior posture, presenting dorsum but neither face nor body outline, represents a considerably more difficult task; and (4) the hidden posture, showing only parts of pelage, represents the most difficult detection task.

**Figure 1.**
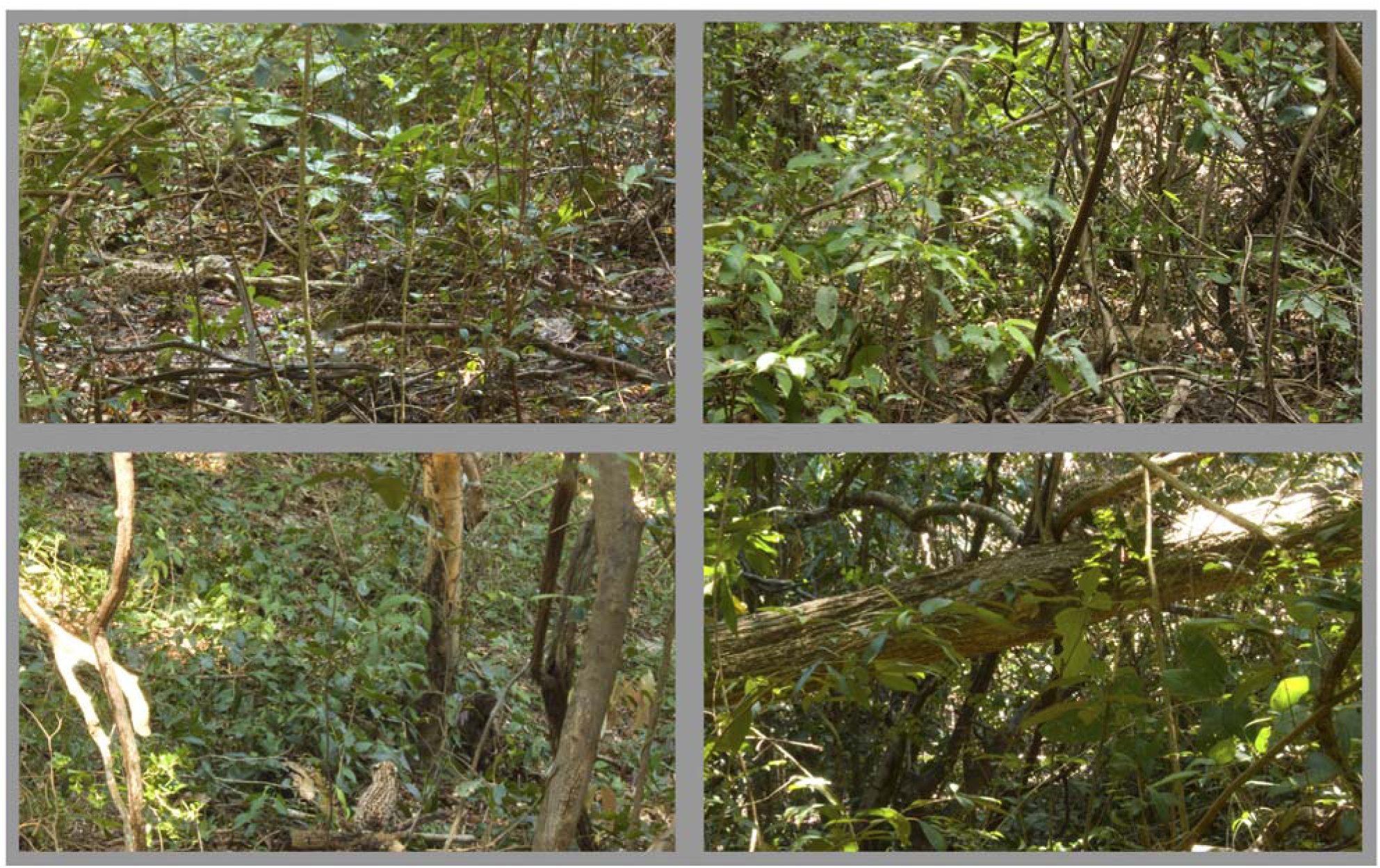
Examples of photographs used as naturalistic stimuli, all of which present a taxidermized model of *Leopardus* sp. in the forest background. A) lateral posture, with the carnivore model positioned in the left side of the scene. B) anterior posture, with the model positioned in the bottom-right area of the scene. C) posterior posture, presenting the model at the bottom of the image. D) hidden posture, where the model is visually obstructed by a trunk at the top of the scene.

We used the Pantone ColorChecker Passport Camera Calibration Software (X-Rite Inc.) to create custom DNG profiles based on colour chart photos, and Adobe Camera RAW to apply these profiles to RAW pictures for colour adjustments. RAW colour adjusted photos were converted to TIFF and edited in Adobe Photoshop CS5. We controlled for real size difference between the carnivore models by creating two stimulus dimension categories, in which the TIFF images were edited so that all models had one of the following dimensions: (1) small targets, in which model’s length (from snout to hindquarter) was set to two centimetres of the monitor screen; and (2) large targets, in which the model’s length was set to five centimetres of the monitor screen. In photographs where specimens were not positioned on their lateral side (i.e., anterior and posterior posture categories), the photograph has been edited so that carnivore model presented the same shoulder height or the same hip height as in its lateral posture photo. These stimulus dimension categories simulate different detection distances: while small specimens appear to be farther away from the observer, larger specimens appear to be closer. We chose these specific stimuli dimensions because they activate retinal area of different sizes when observed on a monitor at a fixed distance of 40 centimetres from the human eye. Under these conditions, small targets stimulate a retinal area smaller than the human fovea, while larger targets stimulate areas that exceed foveal space [77].

### Experimental procedure

We carried out the experiments in the Laboratory of Human Behaviour Evolution of Universidade Federal do Rio Grande do Norte, in individual sessions. The experimental subjects sat in front of a TouchSmart IQ510br 22” touch-screen monitor (Hewlett Packard) and kept their heads at a distance of 40 cm from the screen with the aid of a chin and forehead support. We calibrated the monitor before each experimental session using a colorimeter (Pantone hueyPRO, X-Rite, Inc.), which remained connected to the computer throughout the experiment to ensure that variations in natural ambient light were compensated for by monitor emission. We used a custom-made stimulus presentation software to quantify (1) the latency and (2) the response accuracy of experimental subjects during the detection tasks of carnivore models. This software performed a training session at the beginning of each experiment to familiarize the subjects with the experimental protocol. Training session used only one carnivore model (a specimen of *Leopardus* sp.) and this model was excluded from experimental session to avoid the creation of a search image by subjects.

Experimental sessions started with the presentation of a grey screen for three seconds, after which a + symbol appeared in the centre of the monitor. Subjects were instructed to touch the + to move to the next stage of the experiment, fixing their eyes on the centre of the grey screen while performing this action. After touching +, a set of four photographs was displayed on the monitor, with one photo in each quadrant. Three of the four photos represented a negative discriminative stimulus (i.e., natural scenes without the carnivore model) and the remaining photo represented the positive discriminative stimulus (i.e., natural scene with the carnivore model). The software displayed the photos for one minute or until subjects responded by touching the screen. We instructed subjects to identify the positive stimulus and touch its corresponding photo as quickly as they could. After the subjects’ response or after one minute, the photo set disappeared and was replaced by the grey screen. After three seconds, the + symbol appeared again in the centre of the monitor and, after being touched, a new experimental trial began. The software presented the grey screen between each experimental trial to minimize the possibility of retinal postimages interfering with stimuli visualization.

We selected a total of 72 photographs to be presented to experimental subjects as positive stimuli. Each photograph represented a unique combination of all categories (three carnivore models x three background scenarios x four body postures x two stimulus dimensions = 72). We doubled the number of positive stimuli by flipping all photos horizontally during image editing, creating a new set of 72 photographs. Thus, the software displayed a total of 144 sets of photographs for all experimental subjects, and each set presented a unique positive stimulus. The order of presentation of each photo set varied randomly between individual experimental sessions, and the position of positive and negative stimuli in the four monitor quadrants varied randomly between each photo set. We believe that this experimental procedure minimized or nullified any observer bias during behavioural sampling, since experiments were conducted and data were recorded by a computer software.

### Statistical analysis

The statistical analyses followed a previous study that employed a similar method [62]. We analysed data with linear mixed models (LMM, response latency) and generalized linear mixed models (GLMM, detection accuracy) [78], including subject identity as a random intercept in all the models to account for the repeated measure design. Here, our primary focus was to examine whether the response of subjects (response latency and detection accuracy) varied with carnivore body posture depending on the visual phenotype (dichromat or trichromat). Therefore, nearly all models include body posture and visual phenotype as fixed effects plus the interaction between these two variables.

We were also interested in evaluating whether body posture affects differentially the responses of dichromats and trichromats according to background scenario (grassland, savanna, or forest), carnivore species (lesser grison, ocelot, or cougar), and stimulus dimension (small or large). To prevent overfitting driven by the excess of correlated explanatory variables [62], we split models and built one model for each level of background scenario (3 levels), carnivore species (3 levels), and stimulus dimension (2 levels), totalling 8 models for each response variable (response latency and detection accuracy). Because subjects’ response varies with background scenario, carnivore species, and stimulus dimension [62], we added two of these variables as covariates when appropriate. For example, we added carnivore species and stimulus dimension as fixed effects in the models built for each level of background scenario.

Response latency was log-transformed (natural log) before the analyses to meet the normality assumption. For the detection accuracy analysis, we analysed whether and how the probability of correct responses (Bernoulli family) varies with visual phenotype, body posture, and covariates. We choose for each model the link function (*logit*, *probit*, or *cloglog*) that generated the lowest value for the second-order Akaike’s information criterion – AICc [78]. Using the same dataset, de Moraes *et al.* [62] found that a ceiling effect driven by the long trial duration (one minute) masked the differences between phenotypes in detection accuracy. Therefore, we computed the detection accuracy of subjects in the first three seconds of each trial as in the previous study.

We also assessed whether facial colour patterns affect gaze detection of different carnivores. For this purpose, we built two models (one for each response latency and detection accuracy) using only data from the anterior posture of taxidermized models. We included visual phenotype, carnivore species, and their interaction term as fixed effects in these two models. We considered background scenario and stimulus dimension as covariates in the models.

Models were built using *glmmTMB* package from R (version 4.1.1). We assessed the significance of explanatory variables and interactions using Type II Wald χ^2^ tests. We performed post-hoc tests on full models or models without non-significant interaction terms using *emmeans* package [79]. Post-hoc comparisons were made across body postures within each visual phenotype and between visual phenotypes within each body posture (Tukey method of p-value adjustments).

## Results

Body posture affected response latency and detection accuracy irrespective of visual phenotype in most experimental conditions (Figs 2-7, Tables S1-S6). Overall, subjects presented the fastest and most accurate responses to predators in the lateral posture and the slowest and least accurate responses to predators in the hidden posture (for remembering body postures, see Fig. 1). However, in most conditions, subjects presented similar detection performance (latency and accuracy) regardless of whether predators were in the anterior or the lateral postures (Figs. 3-7, Tables S2-S6). Also, in general, subjects responded faster and more accurately to predators in the anterior posture than to predators in the posterior posture. These results were consistent across background scenarios, carnivore species, and stimulus dimensions with some exceptions (Figs. 2-7, Tables S1-S6). A noteworthy exception to this pattern was observed during ocelot detection, in which body posture did not affect response latency except when those carnivores were hidden (which led to slower responses) (Fig. 3, Table S2).

**Figure 2.**
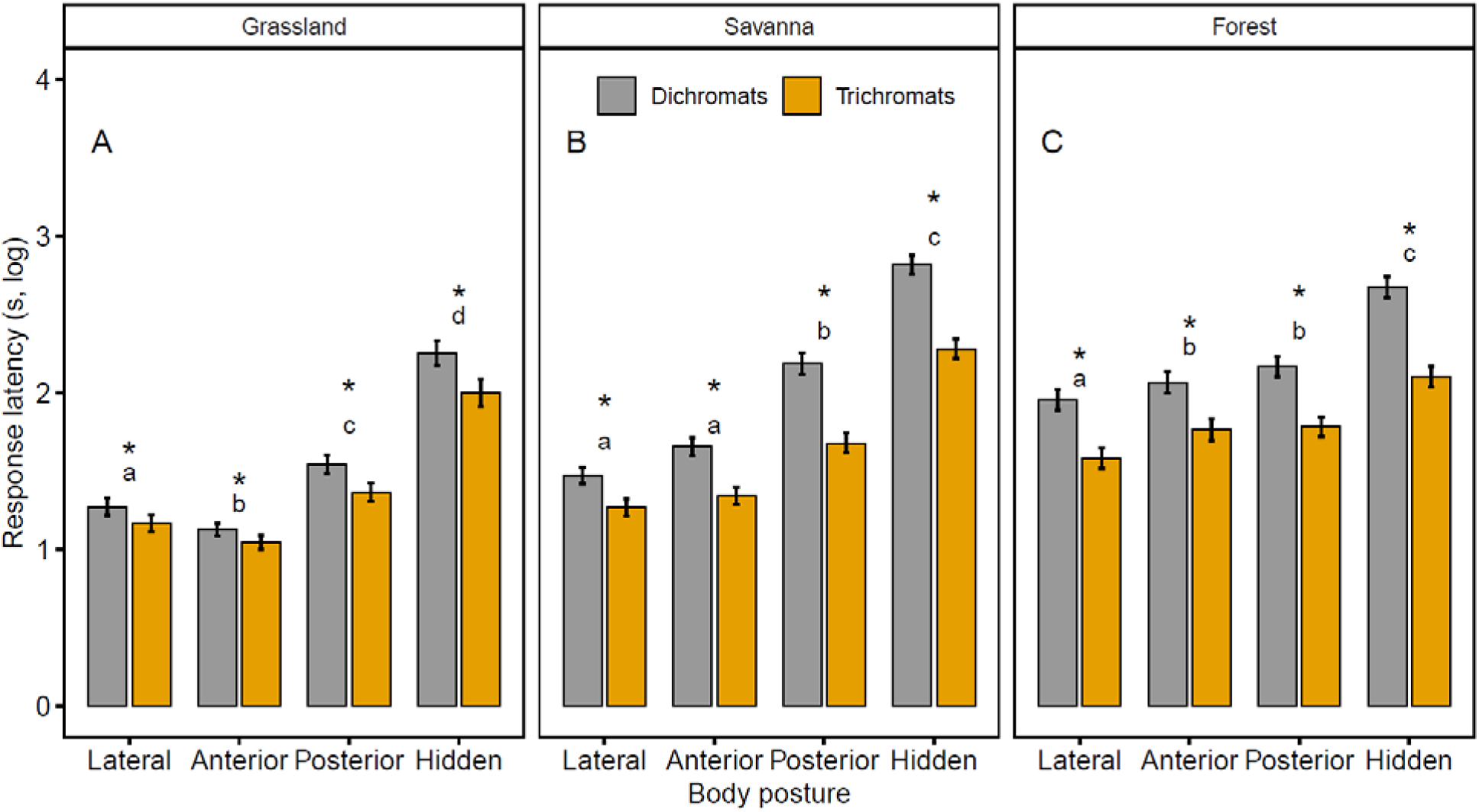
Latency (mean ± se) of trichromats and dichromats to detect taxidermized carnivores in four different body postures and three background scenarios. Differences in response latency between visual phenotypes are highlighted by asterisks. Differences in response latency between body postures are given by different letters.

**Figure 3.**
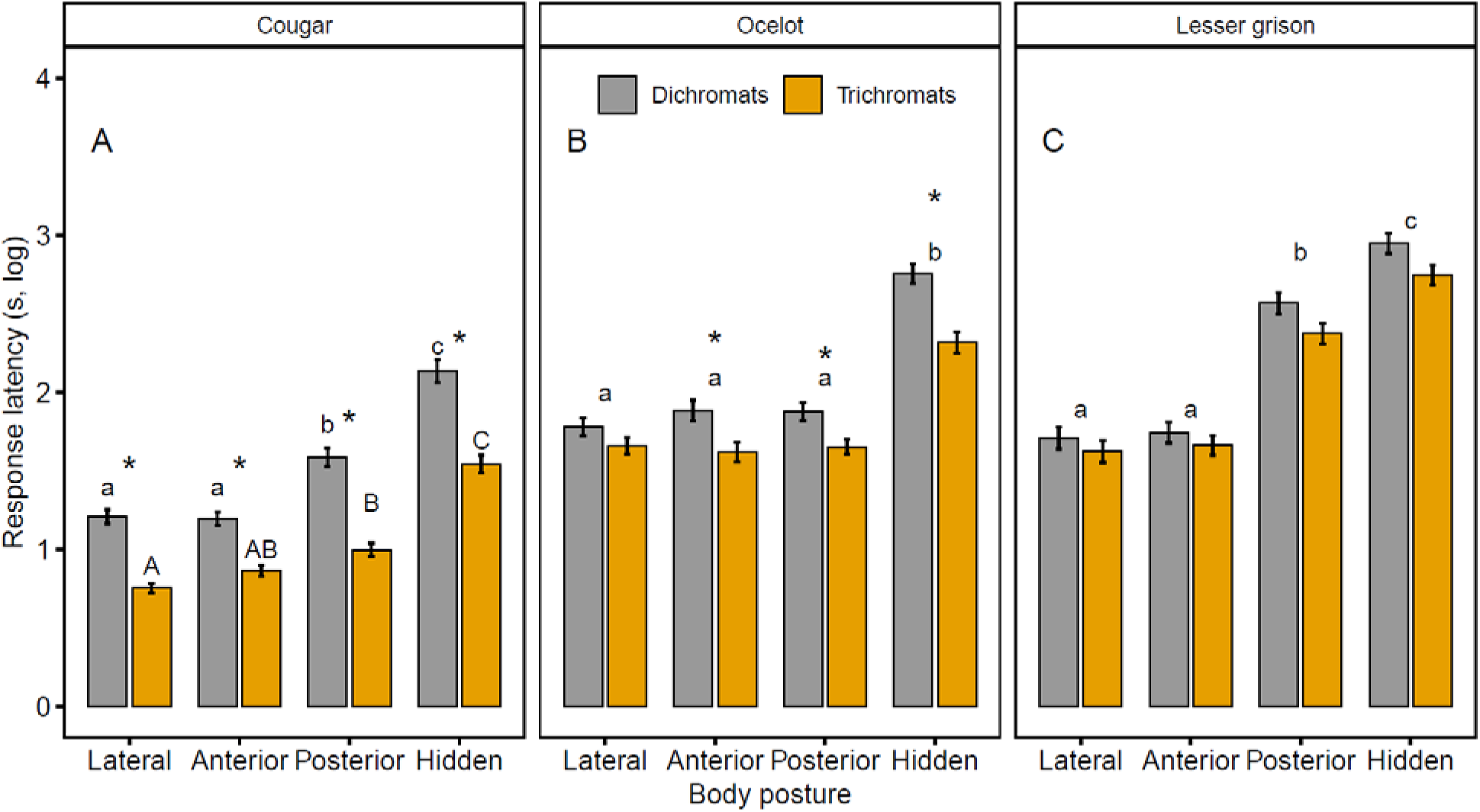
Latency (mean ± se) of trichromats and dichromats to detect three taxidermized carnivores in four different body postures. Differences in response latency between visual phenotypes are highlighted by asterisks. Differences in response latency between body postures are given by different letters.

Irrespective of body posture (Figs 2-7, Tables S1-S6), trichromats outperformed dichromats when detecting cougars, but both visual phenotypes performed similarly when detecting lesser grisons. Ocelots were equally easy to detect by both visual phenotypes but only when positioned laterally; under the remaining postures, trichromats outperformed dichromats (Fig. 3, Table S2). In several conditions, dichromats, but not trichromats, performed worse when detecting predators in the posterior posture than in the anterior posture (Figs. 2, 3, and 7, Tables S1-S2, S6).

### Gaze detection of carnivores

Both trichromats and dichromats performed better in detecting the gaze of cougars (latency and accuracy) than the gaze of ocelots and lesser grisons (Fig. 8, Tables S7-S8). Trichromats outperformed dichromats in detecting the gaze of cougars (latency and accuracy) and ocelots (latency), but these phenotypes did not differ in the ability to detect the gaze of lesser grisons (Fig. 8, Tables S7-S8).

## Discussion

Our results show a broad trichromatic advantage for the detection of carnivoran mammals. This advantage occurred regardless of the body posture presented by animal models and even in some particularly difficult tasks, such as the detection of hidden targets in the most complex and low-light scenario (i.e., forest; Figs. 2.C and 5.C). Trichromats were more efficient than dichromats in detecting potential predators in all background scenarios tested (Figs. 2 and 5), during the detection of orange or yellowish targets (i.e., cougars and ocelots; Figs. 3 and 6), when targets were positioned far from the observer (i.e., small targets; Figs. 4.B and 7.B), and, especially, when animal models showed no relevant visual cues such as body contour or gaze (i.e., posterior and hidden postures, Figs. 2-7). Trichromats and dichromats presented similar performances only in some of the easier tasks (e.g., detection of larger targets with visible body outline or gaze; Fig. 7.A) and during lesser grison detection, the only carnivore model with cryptic coat (i.e., the hardest detection tasks) (Figs. 3.C and 6.C).

**Figure 4.**
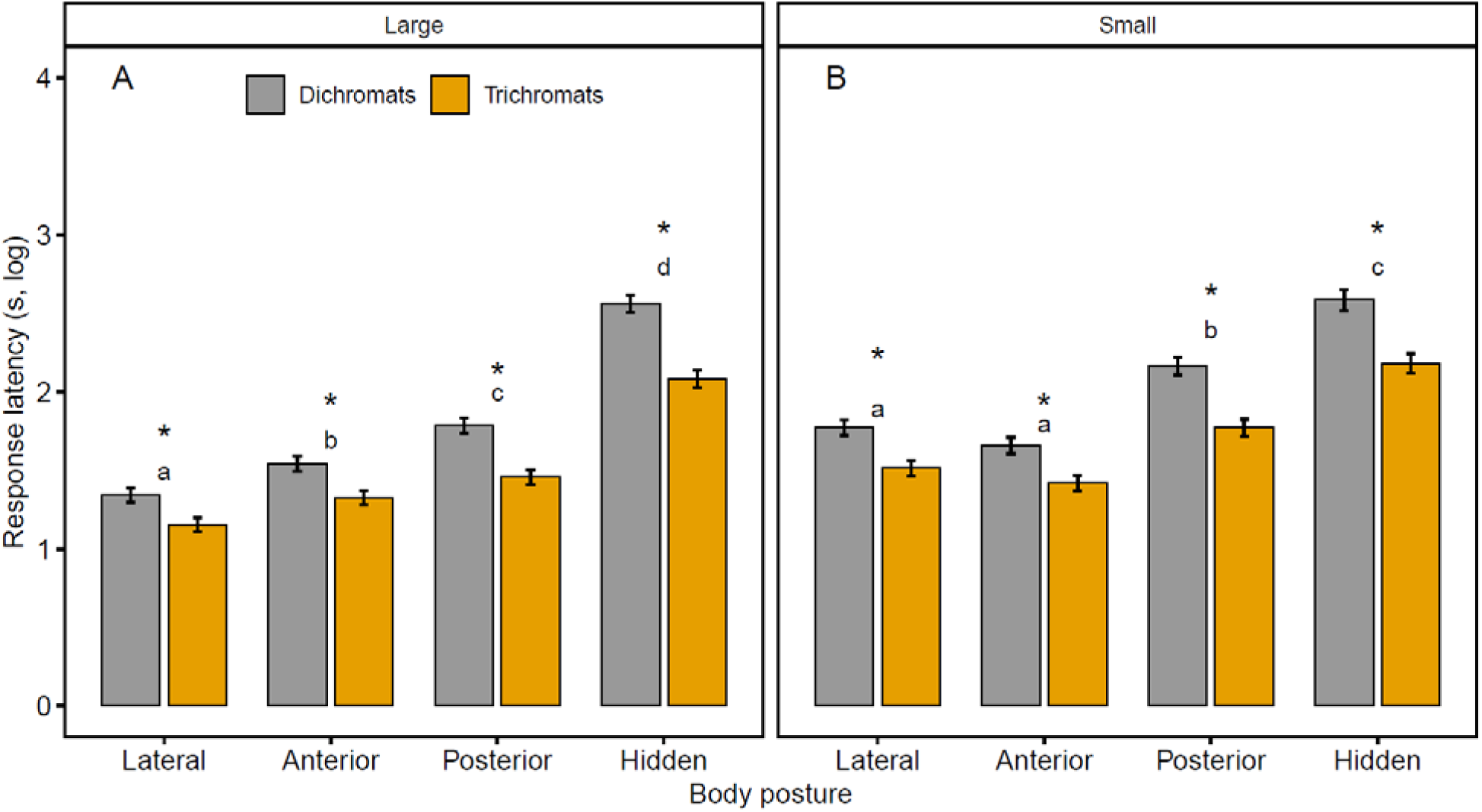
Latency (mean ± se) of trichromats and dichromats to detect taxidermized carnivores in four different body postures and two dimensions. Differences in response latency between visual phenotypes are highlighted by asterisks. Differences in response latency between body postures are given by different letters of the same size.

**Figure 5.**
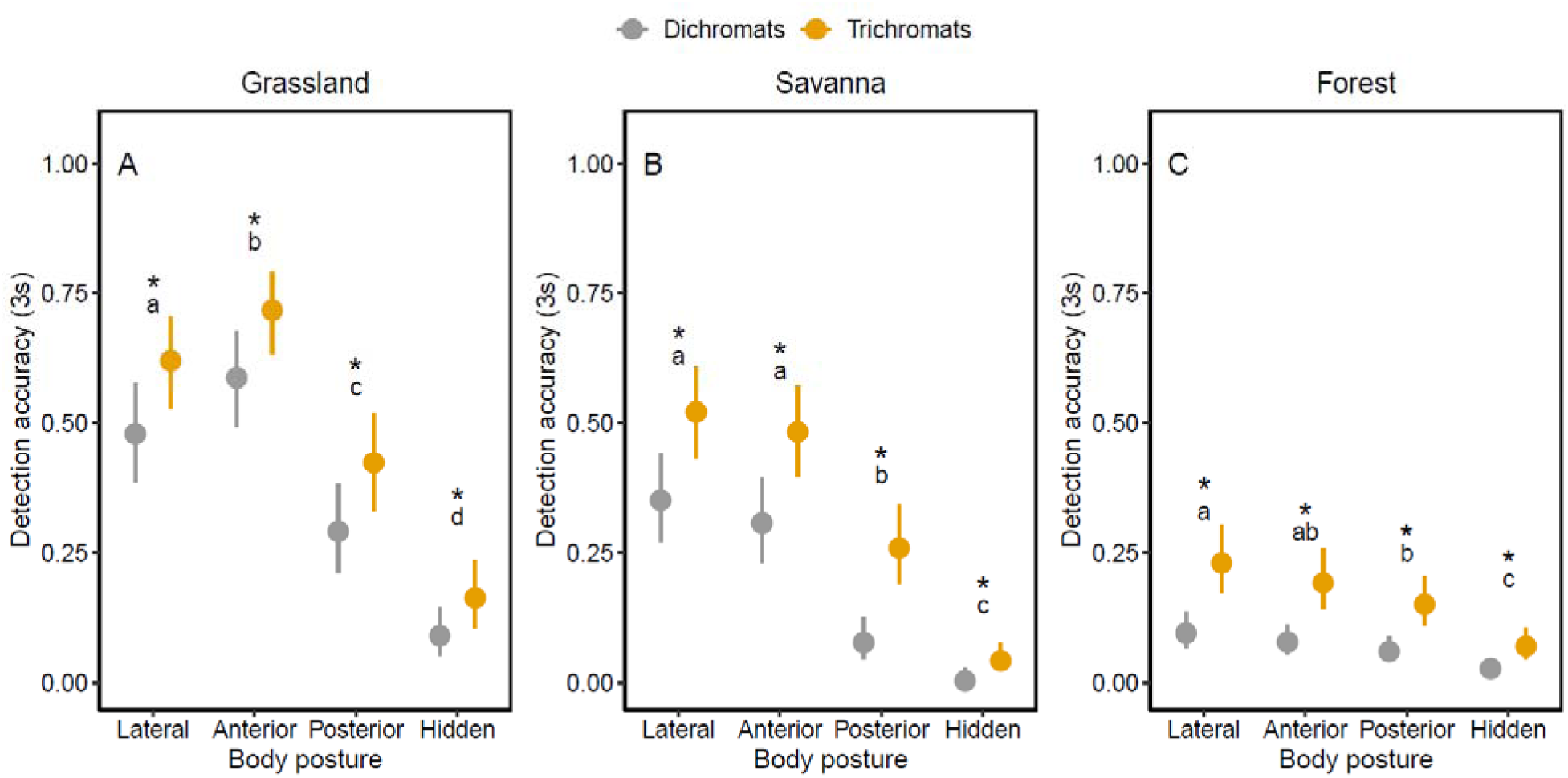
Accuracy of trichromats and dichromats to detect taxidermized carnivores in four different body postures and three background scenarios during the first three seconds of each trial. Detection accuracy consists in model predicted values (mean and 95% CI). Differences in detection accuracy between visual phenotypes are highlighted by asterisks. Differences in detection accuracy between body postures are given by different letters.

**Figure 6.**
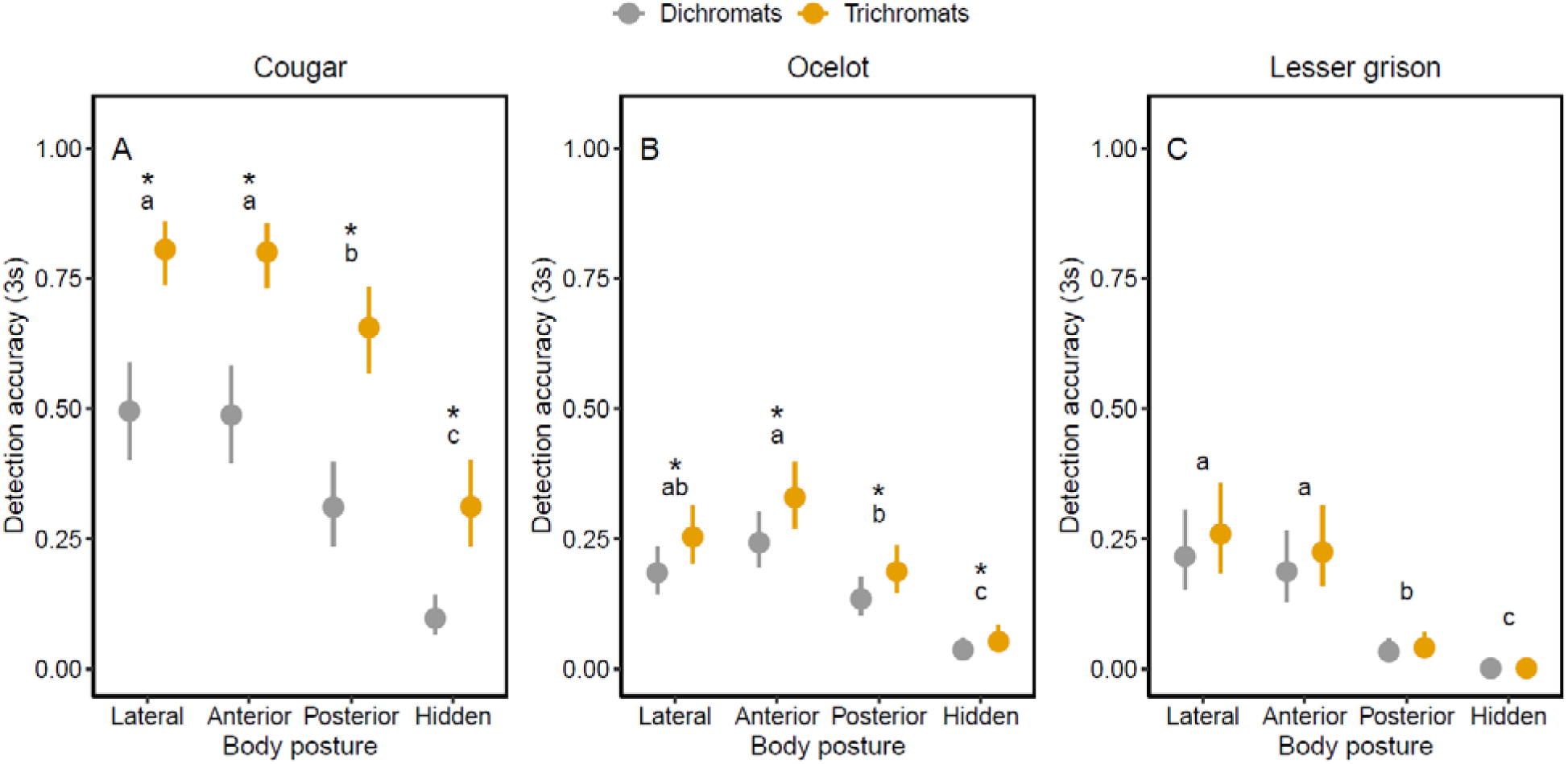
Accuracy of trichromats and dichromats to detect three taxidermized carnivores in four different body postures during the first three seconds of each trial. Detection accuracy consists in model predicted values (mean and 95% CI). Differences in detection accuracy between visual phenotypes are highlighted by asterisks. Differences in detection accuracy between body postures are given by different letters.

The first study to observe this trichromatic advantage on predator detection was conducted by Pessoa *et al.* [61] through visual modelling and behavioural experiments. They confirmed that trichromats not only have a more sensitive sensory apparatus for carnivore mammal perception, but they also detect these targets more easily than dichromats. De Moraes *et al.* [62] took a step further by evaluating in which contexts this trichromatic advantage may be more or less relevant, such as in different background scenarios, during the detection of predators with different colour patterns or at different distances from the observer. The present study goes even further by considering that carnivoran mammals do not fully present themselves to their prey, but instead try to make their own detection difficult by assuming stealthy postures during stalking behaviour [4,6,7]. By evaluating the influence of predator body posture, we observed that trichromacy is even more relevant in facilitating the identification of carnivoran mammals in contexts of enhanced detection difficulty. These results indicate that trichromats are probably more able to thwart attack attempts made by carnivoran mammals, which should increase the survivability of this visual phenotype in comparison to dichromats.

Our results also illustrate a clear process of evolutionary arms race [80] between prey sensory apparatus and predator camouflage capabilities. If we consider an ancestral population of naïve prey that does not know a new predator, we can expect that the latter is not interpreted as a threat [81–83]. Therefore, this new predator probably does not need stealthy behaviours to capture naïve prey. However, if this prey population is visually oriented, natural selection should favour those that visually recognize the predator and move away from it before the attack is carried out [6]. Our results indicate that visual recognition of potential predators is a well-established ability, since experimental subjects easily encountered animals that presented their complete body contour (i.e., lateral posture) during detection tasks (Figs. 2-7). With the evolution of prey’s ability to visually recognize predator’s silhouette, natural selection should favour, in response, those predators that visually obstruct their bodies while hunting [6]. This scenario probably favoured the evolution of stalking behaviour by making the presence of predators less obvious for their prey. However, even if these ancient predators manage to occlude parts of their bodies, they still need to visually monitor prey to carry out the attack, leaving their gaze available as an environmental cue [5,84].

**Figure 7.**
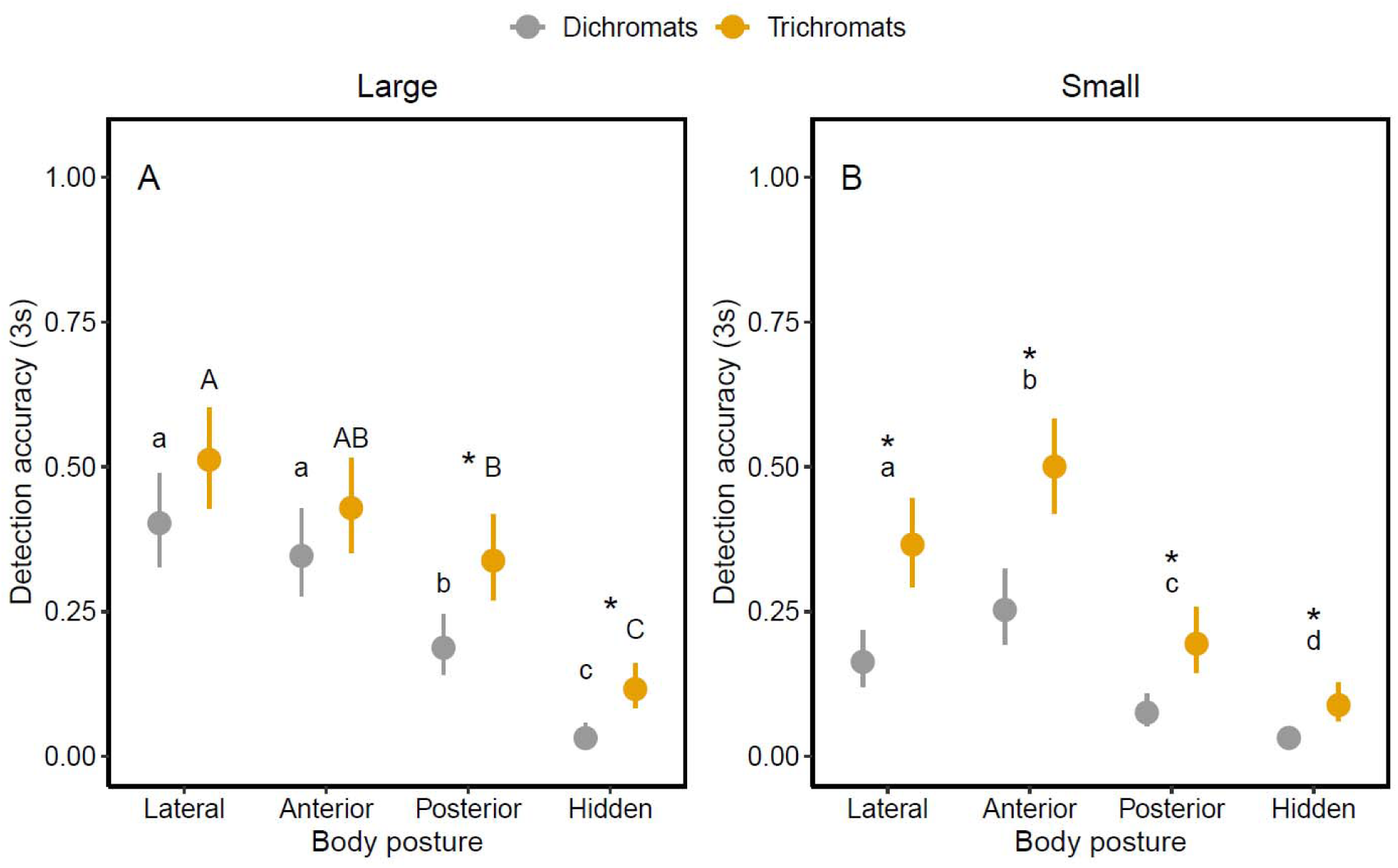
Accuracy of trichromats and dichromats to detect taxidermized carnivores in four different body postures and two dimensions during the first three seconds of each trial. Detection accuracy consists in model predicted values (mean and 95% CI). Differences in detection accuracy between visual phenotypes are highlighted by asterisks. Differences in detection accuracy between body postures are given by different letters of the same size.

The display of predator’s gaze can generate a new evolutionary pressure on prey, in which natural selection should favour those that have the neural capacity to infer a higher predation risk from the visual cue represented by two staring eyes [5]. Our results show that, on many occasions, experimental subjects presented similar performances while detecting laterally or frontally positioned targets (e.g., Figs. 2.B and 5.B), with one instance where subjects found targets in the frontal posture more easily than targets in the lateral posture (Figs. 2.A and 5.A). It was expected that lateral posture would require shorter detection latency and result in higher detection accuracy than the other postures since it presents the animal’s complete body contour, but also because lateral positioned models represent the largest stimuli (i.e., they activate a larger retinal area [67,68]). However, images presenting predators’ faces (i.e., frontal posture) can trigger similar or superior behavioural responses to those generated by larger stimuli (i.e., lateral posture), even though they activate a comparatively smaller retinal area. These results indicate that prey can readily perceive a predator even if its body is not clearly visible in the environment and perform an adequate antipredator response based on the interpretation of two staring eyes as a high predation risk cue. Therefore, two distinct search images can be equally relevant in predator detection tasks: the predator’s body contour and its gaze. These results also point to the importance of gaze recognition for primates (at least for humans), indicating that predation risk may have joined social communication pressures and influenced the evolution of the mental ability to detect and extract relevant information from eyes in the environment [18,85].

A probable next step in this sensory arms race is the evolution of gaze camouflage. As prey are able to infer high predation risk from staring eyes, natural selection may have favoured predators that add noise to the visual perception of their faces [14,28]. This noise likely hampers gaze identification, allowing predators to get even closer to prey during stalking behaviour and achieve higher capture success [8]. Our results seem to corroborate this reasoning, considering that the predator that presents facial uniform colouration (i.e., cougar) is more easily detected than models that present complex facial colourations (i.e., ocelots and lesser grison) (Fig. 8). Caro *et al.* [28] investigated the evolution of anterior colouration of carnivoran mammals through phylogenetic analyses, and they found that the anterior colour patterns of families Viverridae and Herpestidae are associated with the proportion of visually oriented prey (i.e., mammals) in the diet, indicating that anterior colouration may function as disruptive camouflage in these two families. However, their analyses failed to find similar results for Felidae and Mustelidae, the two families with animal models used in the present study. According to Caro *et al.* [28], the anterior colouration of mustelids is associated with the ability to spray noxious anal secretions and with pugnacious behaviour, while anterior colouration of felids presented no relationship with any investigated variable. Our results, nevertheless, are clear in showing that the complex facial colouration presented by a mustelid (lesser grison) and some felids (ocelots) hinder anterior recognition of these animals, in comparison with anterior uniform colouration of another felid (cougar). Thus, the complex facial colour patterns of mustelids and felids may also have evolved by camouflaging the gaze of these carnivores to their prey, adding to intra and interspecific communication as possible factors that influenced the evolution of facial colouration in Carnivora [28].

**Figure 8.**
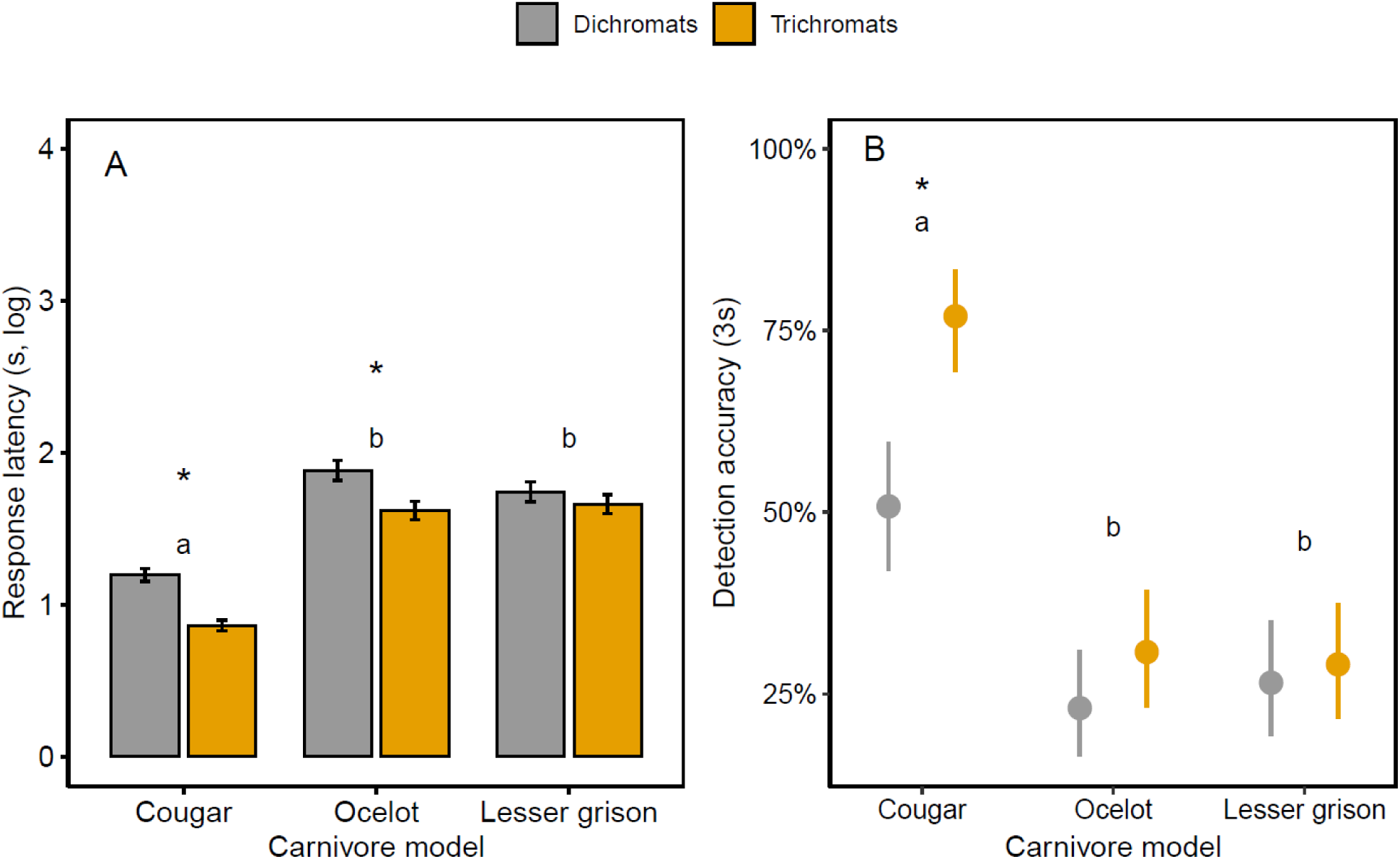
Latency (mean ± se) and accuracy of trichromats and dichromats to detect taxidermized carnivores showing their faces and eyes (i.e., anterior posture). Detection accuracy consists in model predicted values during the first three seconds of each trial (mean and 95% CI). Differences in detection accuracy between visual phenotypes are highlighted by asterisks. Differences in detection accuracy between carnivore models are given by different letters.

An important step in the arms race described here is represented by the evolution of trichromacy. Although this visual phenotype probably evolved in feeding contexts, such as the detection of reddish or orange fruits, leaves and other food items [40,42,43,46,50–52,86], it is possible that an exaptation process [87] has occurred once trichromacy helped some primates to more easily identify their predators. Thus, as proposed by de Moraes *et al.* [62], predation risk may be one of the selective forces that maintain trichromacy in current primates, especially in the parvorder Catarrhini (i.e., the uniformly trichromats primates). Our results, in general, demonstrate the superiority of trichromacy in this context, but some specific situations make even clearer how this visual phenotype was especially important in the sensory arms race between primates and their mammalian predators. Under some experimental conditions, trichromats showed similar performances when detecting anteriorly or posteriorly positioned predators (Figs. 3.A, 3.B, and 7.A), a response not observed in dichromats. It is probable that trichromats have the ability of developing a third search image for predator detection tasks: coat colour. In line with this, Coss *et al.* [5] demonstrated that wild bonnet macaques (*Macaca radiata*) recognized spotted yellow morphs of leopards (*Panthera pardus*) faster than the rare dark melanic forms, suggesting that this catarrhine uses leopard’s most common colour pattern as a cue for predator identification. The yellowish/orange coat presented by some carnivoran mammals possibly evolved by camouflaging these predators to their prey, which are usually other mammals with a dichromatic visual phenotype [33]. Therefore, the coat colour of some carnivores represents a search image for trichromats that is not available to dichromats, since this visual phenotype does not present the necessary sensory channels to perceive this environmental cue [61]. This trichromatic advantage is also observed when we evaluate the effect of gaze camouflage (Fig. 8). Although complex colour patterns hampered trichromats’ performance during predators’ gaze detection, in comparison to uniform coloured faces, trichromats performed better than dichromats when targets presented complex orange/yellowish facial colours.

The colour pattern presented by lesser grison seems to be immune to trichromacy, since the performances of trichromats and dichromats were similar during the detection of this carnivore in all evaluated body postures (Figs. 3 and 6). The lesser grison’s facial colour pattern is also able to nullify the effect of trichromacy on gaze detection (Fig. 8). Therefore, the cryptic colours presented by this carnivore appear to be the final (or contemporary) step in the sensory arms race described so far, as it allows these animals to overcome the evolutionary novelty presented by primates (i.e., trichromacy) and effectively camouflage themselves against dichromatic and trichromatic prey. Lesser grison are not primate predators [74] and probably were not involved in any recent evolutionary interaction with primates. However, it is important that future studies investigate whether primate specialist predators (or those that prey upon any other trichromatic animal) tend to show coat (or plumage or scales) colour pattern in shades outside the yellowish/orange visual spectrum.

In conclusion, we demonstrated the efficiency of primate trichromacy in predator detection tasks, this time considering the influence of postures performed by these same predators while stalking their prey. Although predators may act stealthy and suppress traditional search images that indicate their presence on the environment (i.e., body contour and gaze), trichromatic prey can easily identify them by using coat colour as a high-risk cue. An important factor not evaluated by our experiments is the influence of movement [88,89] in carnivore detection. The hunting behaviour of carnivoran mammals is a succession of static and moving moments [4], so that our experiments only contemplated the static stages. Experiments that assess whether a predator’s colour pattern difficult its identification by prey during movement would be very informative. Specifically, it would be interesting to compare the performances of different camouflage types (e.g., background matching with uniform colouration, or background matching with complex patterns of spots and stripes, or disruptive colouration, etc.) on the detection of moving predators.

Going further, our results point to a comprehensive advantage of trichromacy in different contexts of background scenery, predators’ coat colour and detection distances, especially in situations of greater detection difficulty. A productive step for future studies is to investigate whether this trichromatic advantage in predator detection also applies to animals from other non-mammalian groups that also prey on primates, such as birds of prey [90] and snakes [91]. It is especially relevant to consider these groups since orange hues are uncommon among avian or snake mammalian predators, which could reduce the importance of trichromacy in these contexts. Therefore, visual modelling studies or behavioural experiments that use birds of prey and snakes as detection targets are essential for understanding the evolution of primate colour vision, either reinforcing the relevance of trichromacy, or meeting a balancing force [92] with a theoretical dichromatic advantage on the camouflage breaking [63,65,66] of cryptic predators. We once again demonstrate that the research of primate colour vision evolution can take multiple paths besides the investigation of relative phenotype advantages in foraging, so we emphasize the importance of considering other selective pressures in experimental studies, such as intraspecific communication (e.g., [59,60,93]) and predation risk.

## Acknowledgements

This study was financed in part by the Coordenacao de Aperfeicoamento de Pessoal de Nivel Superior – Brasil CAPES (Finance Codes 001 and 043/2012), by Conselho Nacional de Desenvolvimento Cientifico e Tecnologico – Brasil (CNPq) (Finance Codes 478222/2006-8 and 474392/2013-9) and by Programa de Apoio aos Nucleos de Excelencia – FAPERN/CNPq (Finance Code 25674/2009). A M.Sc. Scholarship was granted to P.Z.M and a Researcher Scholarship was granted to D.M.A.P. both by Conselho Nacional de Desenvolvimento Cientifico e Tecnologico – Brasil CNPq. P.D. received a post-doctoral fellowship from CAPES (88887.469218/2019-00).

