## Supplementary material for "Body posture, gaze, and predator detection: how stalking behaviour may influence colour vision evolution": Table

Table S1. Type II Wald χ^2^ tests in linear mixed models showing the effects of visual phenotype and stimulus position in the detection latency (log) for each background scenario while accounting for carnivore species and stimulus dimension. df = degrees of freedom.

|  | Grassland^a^ | | Savanna^b^ | | Forest^c^ | |
| --- | --- | --- | --- | --- | --- | --- |
| Predictor (df) | χ^2^ | P | χ^2^ | P | χ^2^ | P |
| Visual phenotype (1) | 4.22 | 0.04 | 26.11 | < 0.0001 | 129.4 | < 0.0001 |
| Stimulus position (3) | 584.48 | < 0.0001 | 564.37 | < 0.0001 | 26.79 | < 0.0001 |
| Carnivore species (2) | 632.10 | < 0.0001 | 286.20 | < 0.0001 | 495.48 | < 0.0001 |
| Stimulus dimension (1) | 47.24 | < 0.0001 | 22.08 | < 0.0001 | 71.81 | < 0.0001 |
| Visual phenotype × Stimulus position (3) | 3.85 | 0.28 | 13.48 | 0.004 | 4.10 | 0.25 |

Random intercept standard deviation: subject identity = 0.21^a^, 0.19^b^, 0.20^c^.

Table S2. Type II Wald χ^2^ tests in linear mixed models showing the effects of visual phenotype and stimulus position in the detection latency (log) for each carnivore species while accounting for background scenario and stimulus dimension. df = degrees of freedom.

|  | Lesser grison^a^ | | Ocelot^b^ | | Cougar^c^ | |
| --- | --- | --- | --- | --- | --- | --- |
| Predictor (df) | χ^2^ | P | χ^2^ | P | χ^2^ | P |
| Visual phenotype (1) | 1.33 | 0.24 | 12.85 | 0.003 | 57.65 | < 0.0001 |
| Stimulus position (3) | 485.65 | < 0.0001 | 261.28 | < 0.0001 | 532.43 | < 0.0001 |
| Background scenario (2) | 225.50 | < 0.0001 | 40.97 | < 0.0001 | 335.03 | < 0.0001 |
| Stimulus dimension (1) | 15.56 | < 0.0001 | 126.40 | < 0.0001 | 22.80 | < 0.0001 |
| Visual phenotype × Stimulus position (3) | 0.70 | 0.87 | 7.99 | 0.046 | 18.07 | 0.0004 |

Random intercept standard deviation: subject identity = 0.28^a^, 0.18^b^, 0.19^c^.

Table S3. Type II Wald χ^2^ tests in linear mixed models showing the effects of visual phenotype and stimulus position in the detection latency (log) for each stimulus dimension while accounting for background scenario and carnivore species. df = degrees of freedom.

|  | Large^a^ | | Small^b^ | |
| --- | --- | --- | --- | --- |
| Predictor (df) | χ^2^ | P | χ^2^ | P |
| Visual phenotype (1) | 19.08 | < 0.0001 | 17.74 | < 0.0001 |
| Stimulus position (3) | 731.80 | < 0.0001 | 393.59 | < 0.0001 |
| Background scenario (2) | 220.25 | < 0.0001 | 226.04 | < 0.0001 |
| Carnivore species (2) | 612.80 | < 0.0001 | 645.90 | < 0.0001 |
| Visual phenotype × Stimulus position (3) | 13.89 | 0.003 | 4.04 | 0.26 |

Random intercept standard deviation: subject identity = 0.19^a^, 0.21^b^.

Table S4. Analyses of deviance (Type II Wald χ^2^ tests) showing the effects of visual phenotype and stimulus position in the detection accuracy for each background scenario while accounting for carnivore species and stimulus dimension in generalized linear mixed models (Binomial residuals). Here we considered the probability of detection during the first three seconds of each trial. df = degrees of freedom.

|  | Grassland^a^ | | Savanna^b^ | | Forest^c^ | |
| --- | --- | --- | --- | --- | --- | --- |
| Predictor (df) | χ^2^ | P | χ^2^ | P | χ^2^ | P |
| Visual phenotype (1) | 5.23 | 0.02 | 18.31 | < 0.0001 | 37.85 | < 0.0001 |
| Stimulus position (3) | 200.58 | < 0.0001 | 158.10 | < 0.0001 | 21.99 | < 0.0001 |
| Carnivore species (2) | 268.08 | < 0.0001 | 175.04 | < 0.0001 | 140.52 | < 0.0001 |
| Stimulus dimension (1) | 11.51 | 0.0007 | 16.55 | < 0.0001 | 20.35 | < 0.0001 |
| Visual phenotype × Stimulus position (3) | 5.47 | 0.14 | 6.64 | 0.08 | 5.32 | 0.15 |

Random intercept standard deviation: subject identity = 0.43^a^, 0.50^b^, 0.46^c^. Link function: *probit*^a^, *logit*^b^. *cloglog*^c^.

Table S5. Analyses of deviance (Type II Wald χ^2^ tests) showing the effects of visual phenotype and stimulus position in the detection accuracy for each carnivore species while accounting for background scenario and stimulus dimension in generalized linear mixed models (Binomial residuals). Here we considered the probability of detection during the first three seconds of each trial. df = degrees of freedom.

|  | Lesser grison^a^ | | Ocelot^b^ | | Cougar^c^ | |
| --- | --- | --- | --- | --- | --- | --- |
| Predictor (df) | χ^2^ | P | χ^2^ | P | χ^2^ | P |
| Visual phenotype (1) | 0.77 | 0.38 | 5.17 | 0.02 | 35.72 | < 0.0001 |
| Stimulus position (3) | 86.95 | < 0.0001 | 62.32 | < 0.0001 | 185.68 | < 0.0001 |
| Background scenario (2) | 101.97 | < 0.0001 | 62.03 | < 0.0001 | 182.66 | < 0.0001 |
| Stimulus dimension (1) | 2.42 | 0.12 | 54.34 | < 0.0001 | 5.68 | 0.017 |
| Visual phenotype × Stimulus position (3) | 1.67 | 0.64 | 5.35 | 0.15 | 4.06 | 0.25 |

Random intercept standard deviation: subject identity = 0.67^a^, 0.30^b^, 0.65^c^. Link function: *cloglog*^a,c^, *logit*^b^.

Table S6. Analyses of deviance (Type II Wald χ^2^ tests) showing the effects of visual phenotype and stimulus position in the detection accuracy for each stimulus dimension while accounting for background scenario and carnivore species in generalized linear mixed models (Binomial residuals). Here we considered the probability of detection during the first three seconds of each trial. df = degrees of freedom.

|  | Large^a^ | | Small^b^ | |
| --- | --- | --- | --- | --- |
| Predictor (df) | χ^2^ | P | χ^2^ | P |
| Visual phenotype (1) | 7.80 | 0.005 | 24.35 | < 0.0001 |
| Stimulus position (3) | 211.82 | < 0.0001 | 177.53 | < 0.0001 |
| Background scenario (2) | 168.53 | < 0.0001 | 188.18 | < 0.0001 |
| Carnivore species (2) | 274.13 | < 0.0001 | 318.83 | < 0.0001 |
| Visual phenotype × Stimulus position (3) | 13.06 | 0.005 | 3.49 | 0.32 |

Random intercept standard deviation: subject identity = 0.44^a^, 0.58^b^. Link function: *cloglog*^a^, *logit*^b^.

Table S7. Type II Wald χ^2^ tests in linear mixed models showing the effects of visual phenotype in the detection latency (log) of the gaze (i.e., anterior posture) of different carnivore species while accounting for background scenario and stimulus dimension. df = degrees of freedom.

| Predictor (df) | χ^2^ | P |
| --- | --- | --- |
| Visual phenotype (1) | 9.74 | 0.002 |
| Carnivore species (2) | 301.01 | < 0.0001 |
| Background scenario (2) | 298.51 | < 0.0001 |
| Stimulus dimension (1) | 10.61 | 0.001 |
| Visual phenotype × Stimulus position (3) | 6.44 | 0.040 |

Random intercept standard deviation: subject identity = 0.19.

Table S8. Analyses of deviance (Type II Wald χ^2^ tests) showing the effects of visual phenotype in the detection accuracy of the gaze (i.e., anterior posture) of different carnivore species while accounting for background scenario and stimulus dimension in generalized linear mixed models (Binomial residuals). Here we considered the probability of detection during the first three seconds of each trial. df = degrees of freedom.

| Predictor (df) | χ^2^ | P |
| --- | --- | --- |
| Visual phenotype (1) | 7.70 | 0.006 |
| Carnivore species (2) | 138.77 | < 0.0001 |
| Background scenario (2) | 175.32 | < 0.0001 |
| Stimulus dimension (1) | 0.76 | 0.38 |
| Visual phenotype × Stimulus position (3) | 13.00 | 0.002 |

Random intercept standard deviation: subject identity = 0.32. Link function: *probit*.

Source code used in R to run the statistical analyses:

```
# Predator detection and primate color vision
# Update: 29 May 2023

# Analyses including the 39 male subjects

# 1. Latency of detection ~ Stimulus position * Visual phenotype (for each Habitat type)
# - grassland
# - savanna
# - forest
# 2. Latency of detection ~ Stimulus position * Visual phenotype (for each Carnivorous species)
# - lesser grison
# - ocelot
# - cougar
# 3. Latency of detection ~ Stimulus position * Visual phenotype (for each Stimulus dimension)
# - dimension: small
# - dimension: large
# 4. Latency of detection ~ Predator * Visual phenotype (for anterior posture only)
# 5. Detection accuracy ~ Stimulus position * Visual phenotype (for each Habitat type)
# - grassland
# - savanna
# - forest
# 6. Detection accuracy ~ Stimulus position * Visual phenotype (for each Carnivorous species)
# - lesser grison
# - ocelot
# - cougar
# 7. Detection accuracy ~ Stimulus position * Visual phenotype (for each Stimulus dimension)
#   - dimension: small
#   - dimension: large
# 8. Detection accuracy (3s) ~ Predator * Visual phenotype (for anterior posture only)

options(scipen=999)

# Load packages
library(glmmTMB); library(openxlsx); library(car); library(performance); library(DHARMa); library(lme4); library (AICcmodavg);
library(emmeans); library(ggplot2); library(ggpubr); library(parameters); library(psych); library(sjPlot); library(ggeffects)

# Load data
d1<-read.xlsx("Dataset.xlsx")

# Processing data
d1$Subject<-factor(d1$Subject)
d1$VisualPhenotype<-factor(d1$VisualPhenotype)
d1$Background<-factor(d1$Background)
d1$Predator<-factor(d1$Predator)
d1$Size<-factor(d1$Size)
d1$StimulusPosition<-factor(d1$StimulusPosition)
d1$Detection<-factor(d1$Detection,levels=c("no","yes"))
d1$Predator <- factor(d1$Predator, levels = c("lesser grison", "ocelot", "cougar"))
d1$Background <- factor(d1$Background, levels = c("grassland", "savanna", "forest"))
d1$StimulusPosition <- factor(d1$StimulusPosition, levels = c("side", "front", "back", "hide"))
d1$Size<-factor(d1$Size,levels=c("large","small"))

# log transforming latency
d1$loglat<-log(d1$Latency)
d1<-droplevels(d1)

# Removing the female subject
d1<-subset(d1,Subject!="13")
d1<-droplevels(d1)

#####
# 1. Latency of detection ~ Stimulus position * Visual phenotype (for each Habitat type)
#####

##### Grassland
d2<-subset(d1,Background=="grassland")
d2<-na.omit(d2)
# GLMM
m1<-glmmTMB(loglat~VisualPhenotype*StimulusPosition+Predator+Size+(1|Subject),data=d2)
# Model validation
check_model(m1)
check_collinearity(m1) # ignored since it was a product of two variables
# Model fitting
Anova(m1)
summary(m1)
model_parameters(m1,standardize = "refit")
# post hoc
m1<-glmmTMB(loglat~VisualPhenotype+StimulusPosition+Predator+Size+(1|Subject),data=d2)
pairs(emmeans(m1, ~StimulusPosition))
pairs(emmeans(m1, ~VisualPhenotype))

##### Savanna
d2<-subset(d1,Background=="savanna")
d2<-na.omit(d2)
# GLMM
m1<-glmmTMB(loglat~StimulusPosition*VisualPhenotype+Predator+Size+(1|Subject),data=d2)
# Model validation
check_model(m1)
check_collinearity(m1) # ignored since it was a product of two variables
# Model fitting
Anova(m1) # 0.0037
summary(m1)
model_parameters(m1,standardize = "refit")
# post hoc
pairs(emmeans(m1, ~StimulusPosition|VisualPhenotype))
pairs(emmeans(m1, ~VisualPhenotype|StimulusPosition))

##### Forest
d2<-subset(d1,Background=="forest")
d2<-na.omit(d2)
# GLMM
m1<-glmmTMB(loglat~StimulusPosition*VisualPhenotype+Predator+Size+(1|Subject),data=d2)
# Model validation
check_model(m1)
check_collinearity(m1) # ignored since it was a product of two variables
# Model fitting
Anova(m1)
summary(m1)
model_parameters(m1,standardize = "refit")
# post hoc
m1<-glmmTMB(loglat~VisualPhenotype+StimulusPosition+Predator+Size+(1|Subject),data=d2)
pairs(emmeans(m1, ~StimulusPosition))
pairs(emmeans(m1, ~VisualPhenotype))

#####
# 2. Latency of detection ~ Stimulus position * Visual phenotype (for each Carnivorous species)
#####

##### Lesser grison
d2<-subset(d1,Predator=="lesser grison")
d2<-na.omit(d2)
# GLMM
m1<-glmmTMB(loglat~StimulusPosition*VisualPhenotype+Background+Size+(1|Subject),data=d2)
# Model validation
check_model(m1)
check_collinearity(m1) # ignored since it was a product of two variables
# Model fitting
Anova(m1) # position
summary(m1)
model_parameters(m1,standardize = "refit")
# post hoc
m1<-glmmTMB(loglat~StimulusPosition+VisualPhenotype+Background+Size+(1|Subject),data=d2)
pairs(emmeans(m1, ~StimulusPosition))

##### Ocelot
d2<-subset(d1,Predator=="ocelot")
d2<-na.omit(d2)
# GLMM
m1<-glmmTMB(loglat~StimulusPosition*VisualPhenotype+Background+Size+(1|Subject),data=d2)
# Model validation
check_model(m1)
check_collinearity(m1) # ignored since it was a product of two variables
# Model fitting
Anova(m1) # ns
summary(m1)
model_parameters(m1,standardize = "refit")
# post hoc
pairs(emmeans(m1, ~StimulusPosition|VisualPhenotype))
pairs(emmeans(m1, ~VisualPhenotype|StimulusPosition))

##### Cougar
d2<-subset(d1,Predator=="cougar")
d2<-na.omit(d2)
# GLMM
m1<-glmmTMB(loglat~StimulusPosition*VisualPhenotype+Background+Size+(1|Subject),data=d2)
# Model validation
check_model(m1)
check_collinearity(m1) # ignored since it was a product of two variables
# Model fitting
Anova(m1) # ns
summary(m1)
model_parameters(m1,standardize = "refit")
# post hoc
pairs(emmeans(m1, ~StimulusPosition|VisualPhenotype))
pairs(emmeans(m1, ~VisualPhenotype|StimulusPosition))

#####
# 3. Latency of detection ~ Stimulus position * Visual phenotype (for each Stimulus dimension)
#####

##### Stimulus dimension: Large
d2<-subset(d1,Size=="large")
d2<-na.omit(d2)
# GLMM
m1<-glmmTMB(loglat~StimulusPosition*VisualPhenotype+Background+Predator+(1|Subject),data=d2)
# Model validation
check_model(m1)
check_collinearity(m1) # ignored since it was a product of two variables
# Model fitting
Anova(m1) # ns
summary(m1)
model_parameters(m1,standardize = "refit")
# post hoc
pairs(emmeans(m1, ~StimulusPosition|VisualPhenotype))
pairs(emmeans(m1, ~VisualPhenotype|StimulusPosition))

##### Stimulus dimension: Small
d2<-subset(d1,Size=="small")
d2<-na.omit(d2)
# GLMM
m1<-glmmTMB(loglat~StimulusPosition*VisualPhenotype+Background+Predator+(1|Subject),data=d2)
# Model validation
check_model(m1)
check_collinearity(m1) # ignored since it was a product of two variables
# Model fitting
Anova(m1) # ns
summary(m1)
model_parameters(m1,standardize = "refit")
# post hoc
m1<-glmmTMB(loglat~StimulusPosition+VisualPhenotype+Background+Predator+(1|Subject),data=d2)
pairs(emmeans(m1, ~StimulusPosition))
pairs(emmeans(m1, ~VisualPhenotype))

#####
# 4. Latency of detection ~ Carnivore species * Visual phenotype (for anterior posture only)
#####

d2<-subset(d1,StimulusPosition=="front")
d2<-na.omit(d2)
# GLMM
m1<-glmmTMB(loglat~Predator*VisualPhenotype+Background+Size+(1|Subject),data=d2)
# Model validation
check_model(m1)
check_collinearity(m1)
# Model fitting
Anova(m1) # ns
summary(m1)
model_parameters(m1,standardize = "refit")
# post hoc
pairs(emmeans(m1, ~Predator|VisualPhenotype))
pairs(emmeans(m1, ~VisualPhenotype|Predator))

#####
# 5. Detection accuracy (3s) ~ Stimulus position * Visual phenotype (for each Habitat type)
#####

##### Grassland
d2<-subset(d1,Background=="grassland")
d2<-na.omit(d2)
# GLMM
m1<-glmmTMB(Detection~StimulusPosition*VisualPhenotype+Predator+Size+(1|Subject),family="binomial"(link=logit),data=d2)
m2<-glmmTMB(Detection~StimulusPosition*VisualPhenotype+Predator+Size+(1|Subject),family="binomial"(link=probit),data=d2)
m3<-glmmTMB(Detection~StimulusPosition*VisualPhenotype+Predator+Size+(1|Subject),family="binomial"(link=cloglog),data=d2)
AICc(m1);AICc(m2);AICc(m3) # m2
# Model validation
check_model(m2)
check_collinearity(m2)
# Model fitting
Anova(m2)
summary(m2)
model_parameters(m2,standardize = "refit")
# post hoc
m2<-glmmTMB(Detection~StimulusPosition+VisualPhenotype+Predator+Size+(1|Subject),family="binomial"(link=probit),data=d2)
pairs(emmeans(m2, ~StimulusPosition))
pairs(emmeans(m2, ~VisualPhenotype))

##### Savanna
d2<-subset(d1,Background=="savanna")
d2<-na.omit(d2)
# GLMM
m1<-glmmTMB(Detection~StimulusPosition*VisualPhenotype+Predator+Size+(1|Subject),family="binomial"(link=logit),data=d2)
m2<-glmmTMB(Detection~StimulusPosition*VisualPhenotype+Predator+Size+(1|Subject),family="binomial"(link=probit),data=d2)
m3<-glmmTMB(Detection~StimulusPosition*VisualPhenotype+Predator+Size+(1|Subject),family="binomial"(link=cloglog),data=d2)
AICc(m1);AICc(m2);AICc(m3) # m3
# Model validation
check_model(m1)
check_collinearity(m1)
# Model fitting
Anova(m1)
summary(m1)
model_parameters(m1,standardize = "refit")
# post hoc
m1<-glmmTMB(Detection~StimulusPosition+VisualPhenotype+Predator+Size+(1|Subject),family="binomial"(link=logit),data=d2)
pairs(emmeans(m1, ~StimulusPosition))
pairs(emmeans(m1, ~VisualPhenotype))

##### Forest
d2<-subset(d1,Background=="forest")
d2<-na.omit(d2)
# GLMM
m1<-glmmTMB(Detection~StimulusPosition*VisualPhenotype+Predator+Size+(1|Subject),family="binomial"(link=logit),data=d2)
m2<-glmmTMB(Detection~StimulusPosition*VisualPhenotype+Predator+Size+(1|Subject),family="binomial"(link=probit),data=d2)
m3<-glmmTMB(Detection~StimulusPosition*VisualPhenotype+Predator+Size+(1|Subject),family="binomial"(link=cloglog),data=d2)
AICc(m1);AICc(m2);AICc(m3) # m3
# Model validation
check_model(m3)
check_collinearity(m3)
# Model fitting
Anova(m3)
summary(m3)
model_parameters(m3,standardize = "refit")
# post hoc
m3<-glmmTMB(Detection~StimulusPosition+VisualPhenotype+Predator+Size+(1|Subject),family="binomial"(link=cloglog),data=d2)
pairs(emmeans(m3, ~StimulusPosition))
pairs(emmeans(m3, ~VisualPhenotype))

#####
# 6. Detection accuracy (3s) ~ Stimulus position * Visual phenotype (for each Carnivorous species)
#####

##### Lesser grison
d2<-subset(d1,Predator=="lesser grison")
d2<-na.omit(d2)
# GLMM
m1<-glmmTMB(Detection~StimulusPosition*VisualPhenotype+Background+Size+(1|Subject),family="binomial"(link=logit),data=d2)
m2<-glmmTMB(Detection~StimulusPosition*VisualPhenotype+Background+Size+(1|Subject),family="binomial"(link=probit),data=d2)
m3<-glmmTMB(Detection~StimulusPosition*VisualPhenotype+Background+Size+(1|Subject),family="binomial"(link=cloglog),data=d2)
AICc(m1);AICc(m2);AICc(m3) # m3
# Model validation
check_model(m3)
check_collinearity(m3)
# Model fitting
Anova(m3)
summary(m3)
model_parameters(m3,standardize = "refit")
# post hoc
m3<-glmmTMB(Detection~StimulusPosition+VisualPhenotype+Background+Size+(1|Subject),family="binomial"(link=cloglog),data=d2)
pairs(emmeans(m3, ~StimulusPosition))

##### Ocelot
d2<-subset(d1,Predator=="ocelot")
d2<-na.omit(d2)
# GLMM
m1<-glmmTMB(Detection~StimulusPosition*VisualPhenotype+Background+Size+(1|Subject),family="binomial"(link=logit),data=d2)
m2<-glmmTMB(Detection~StimulusPosition*VisualPhenotype+Background+Size+(1|Subject),family="binomial"(link=probit),data=d2)
m3<-glmmTMB(Detection~StimulusPosition*VisualPhenotype+Background+Size+(1|Subject),family="binomial"(link=cloglog),data=d2)
AICc(m1);AICc(m2);AICc(m3) # m3
# Model validation
check_model(m3)
check_collinearity(m3)
# Model fitting
Anova(m3)
summary(m3)
model_parameters(m3,standardize = "refit")
# post hoc
m3<-glmmTMB(Detection~StimulusPosition+VisualPhenotype+Background+Size+(1|Subject),family="binomial"(link=cloglog),data=d2)
pairs(emmeans(m3, ~StimulusPosition))
pairs(emmeans(m3, ~VisualPhenotype))

##### Cougar
d2<-subset(d1,Predator=="cougar")
d2<-na.omit(d2)
# GLMM
m1<-glmmTMB(Detection~StimulusPosition*VisualPhenotype+Background+Size+(1|Subject),family="binomial"(link=logit),data=d2)
m2<-glmmTMB(Detection~StimulusPosition*VisualPhenotype+Background+Size+(1|Subject),family="binomial"(link=probit),data=d2)
m3<-glmmTMB(Detection~StimulusPosition*VisualPhenotype+Background+Size+(1|Subject),family="binomial"(link=cloglog),data=d2)
AICc(m1);AICc(m2);AICc(m3) # m1
# Model validation
check_model(m1)
check_collinearity(m1)
# Model fitting
Anova(m1)
drop1(m1,test=c("Chisq"))
summary(m1)
model_parameters(m1,standardize = "refit")
# post hoc
m1<-glmmTMB(Detection~StimulusPosition+VisualPhenotype+Background+Size+(1|Subject),family="binomial"(link=logit),data=d2)
pairs(emmeans(m1, ~StimulusPosition))
pairs(emmeans(m1, ~VisualPhenotype))

#####
# 7. Detection accuracy (3s) ~ Stimulus position * Visual phenotype (for each Stimulus dimension)
#####

##### Dimension: Large
d2<-subset(d1,Size=="large")
d2<-na.omit(d2)
# GLMM
m1<-glmmTMB(Detection~StimulusPosition*VisualPhenotype+Background+Predator+(1|Subject),family="binomial"(link=logit),data=d2)
m2<-glmmTMB(Detection~StimulusPosition*VisualPhenotype+Background+Predator+(1|Subject),family="binomial"(link=probit),data=d2)
m3<-glmmTMB(Detection~StimulusPosition*VisualPhenotype+Background+Predator+(1|Subject),family="binomial"(link=cloglog),data=d2)
AICc(m1);AICc(m2);AICc(m3) # m3
# Model validation
check_model(m3)
check_collinearity(m3)
# Model fitting
Anova(m3)
summary(m3)
model_parameters(m3,standardize = "refit")
# post hoc
pairs(emmeans(m3, ~StimulusPosition|VisualPhenotype))
pairs(emmeans(m3, ~VisualPhenotype|StimulusPosition))

##### Dimension: Small
d2<-subset(d1,Size=="small")
d2<-na.omit(d2)
# GLMM
m1<-glmmTMB(Detection~StimulusPosition*VisualPhenotype+Background+Predator+(1|Subject),family="binomial"(link=logit),data=d2)
m2<-glmmTMB(Detection~StimulusPosition*VisualPhenotype+Background+Predator+(1|Subject),family="binomial"(link=probit),data=d2)
m3<-glmmTMB(Detection~StimulusPosition*VisualPhenotype+Background+Predator+(1|Subject),family="binomial"(link=cloglog),data=d2)
AICc(m1);AICc(m2);AICc(m3) # m2
# Model validation
check_model(m2); check_collinearity(m2)
# Model fitting
Anova(m1)
summary(m1)
model_parameters(m1,standardize = "refit")
# post hoc
m1<-glmmTMB(Detection~StimulusPosition+VisualPhenotype+Background+Predator+(1|Subject),family="binomial"(link=logit),data=d2)
pairs(emmeans(m1, ~StimulusPosition))
pairs(emmeans(m1, ~VisualPhenotype))

#####
# 8. Detection accuracy (3s) ~ Carnivore species * Visual phenotype (for anterior posture only)
#####

d2<-subset(d1,StimulusPosition=="front")
d2<-na.omit(d2)
d2<-droplevels(d2)
# GLMM
m1<-glmmTMB(Detection~Predator*VisualPhenotype+Background+Size+(1|Subject),family="binomial"(link=logit),data=d2)
m2<-glmmTMB(Detection~Predator*VisualPhenotype+Background+Size+(1|Subject),family="binomial"(link=probit),data=d2)
m3<-glmmTMB(Detection~Predator*VisualPhenotype+Background+Size+(1|Subject),family="binomial"(link=cloglog),data=d2)
AICc(m1);AICc(m2);AICc(m3) # m2
# Model validation
check_model(m2)
check_collinearity(m2)
# Model fitting
Anova(m2)
summary(m2)
model_parameters(m2,standardize = "refit")
# post hoc
pairs(emmeans(m2, ~Predator|VisualPhenotype))
pairs(emmeans(m2, ~VisualPhenotype|Predator))

#####
# Figure 1 - Latency x Background scenario
#####

df<-na.omit(d1)
levels(df$Background)
levels(df$StimulusPosition)
levels(df$StimulusPosition) <- c("Lateral","Anterior","Posterior","Hidden")
levels(df$VisualPhenotype)<-c("Dichromats","Trichromats")
cbPalette <- c("#999999", "#E69F00")
pdf("Fig. 1 - Latency x Background scenario.pdf",width=9,height=5)

{p<-ggbarplot(df,x="StimulusPosition", y = "loglat", add=c("mean_se"),facet.by = c("Background"),
              panel.labs=list(Background=c("Grassland","Savanna","Forest")),color = "black", palette = cbPalette,
              fill="VisualPhenotype",size=0.4,ylim=c(0,4), position = position_dodge(0.8)
)
  p<-p+labs(y="Response latency (s, log)",x="Body posture")
  p<-p+theme(legend.title = element_blank())
  p<-p+theme(strip.background =element_rect(fill="white"))
  p<-p+theme(legend.position = c(0.5, 0.93),legend.direction = "horizontal")
  df$Background<-factor(df$Background)
  len <- length(levels(df$Background))
  vars <- data.frame(expand.grid(levels(df$Background)))
  colnames(vars) <- c("Background")

  # Grassland
  dat <- data.frame(x = rep(1, len), y = rep(1, len), vars, labs=c("a","","")) # side dic
  dat[1, 1:2] <- c(1, 1.5); p<-p + geom_text(aes(x, y, label=labs, group=NULL),data=dat)
  dat <- data.frame(x = rep(1, len), y = rep(1, len), vars, labs=c("b","","")) # front dic
  dat[1, 1:2] <- c(2, 1.3); p<-p + geom_text(aes(x, y, label=labs, group=NULL),data=dat)
  dat <- data.frame(x = rep(1, len), y = rep(1, len), vars, labs=c("c","","")) # back dic
  dat[1, 1:2] <- c(3, 1.8); p<-p + geom_text(aes(x, y, label=labs, group=NULL),data=dat)
  dat <- data.frame(x = rep(1, len), y = rep(1, len), vars, labs=c("d","","")) # hide dic
  dat[1, 1:2] <- c(4, 2.5); p<-p + geom_text(aes(x, y, label=labs, group=NULL),data=dat)
  dat <- data.frame(x = rep(1, len), y = rep(1, len), vars, labs=c("*","",""))
  dat[1, 1:2] <- c(1,1.6); p<-p + geom_text(aes(x, y, label=labs, group=NULL),data=dat,size=6)
  dat <- data.frame(x = rep(1, len), y = rep(1, len), vars, labs=c("*","",""))
  dat[1, 1:2] <- c(2,1.4); p<-p + geom_text(aes(x, y, label=labs, group=NULL),data=dat,size=6)
  dat <- data.frame(x = rep(1, len), y = rep(1, len), vars, labs=c("*","",""))
  dat[1, 1:2] <- c(3,2); p<-p + geom_text(aes(x, y, label=labs, group=NULL),data=dat,size=6)
  dat <- data.frame(x = rep(1, len), y = rep(1, len), vars, labs=c("*","",""))
  dat[1, 1:2] <- c(4,2.6); p<-p + geom_text(aes(x, y, label=labs, group=NULL),data=dat,size=6)
  dat <- data.frame(x = rep(1, len), y = rep(1, len), vars, labs=c("A","",""))
  dat[1, 1:2] <- c(0.75,3.6); p<-p + geom_text(aes(x, y, label=labs, group=NULL),data=dat,size=5)
  p

  # Savanna
  dat <- data.frame(x = rep(1, len), y = rep(1, len), vars, labs=c("","a","")) # side dic
  dat[2, 1:2] <- c(1, 1.7); p<-p + geom_text(aes(x, y, label=labs, group=NULL),data=dat)
  dat <- data.frame(x = rep(1, len), y = rep(1, len), vars, labs=c("","a","")) # front dic
  dat[2, 1:2] <- c(2,1.8); p<-p + geom_text(aes(x, y, label=labs, group=NULL),data=dat)
  dat <- data.frame(x = rep(1, len), y = rep(1, len), vars, labs=c("","b","")) # back dic
  dat[2, 1:2] <- c(3, 2.4); p<-p + geom_text(aes(x, y, label=labs, group=NULL),data=dat)
  dat <- data.frame(x = rep(1, len), y = rep(1, len), vars, labs=c("","c","")) # hide dic
  dat[2, 1:2] <- c(4, 3); p<-p + geom_text(aes(x, y, label=labs, group=NULL),data=dat)
  dat <- data.frame(x = rep(1, len), y = rep(1, len), vars, labs=c("","*",""))
  dat[2, 1:2] <- c(1,1.9); p<-p + geom_text(aes(x, y, label=labs, group=NULL),data=dat,size=6)
  dat <- data.frame(x = rep(1, len), y = rep(1, len), vars, labs=c("","*",""))
  dat[2, 1:2] <- c(2,2); p<-p + geom_text(aes(x, y, label=labs, group=NULL),data=dat,size=6)
  dat <- data.frame(x = rep(1, len), y = rep(1, len), vars, labs=c("","*",""))
  dat[2, 1:2] <- c(3,2.6); p<-p + geom_text(aes(x, y, label=labs, group=NULL),data=dat,size=6)
  dat <- data.frame(x = rep(1, len), y = rep(1, len), vars, labs=c("","*",""))
  dat[2, 1:2] <- c(4,3.2); p<-p + geom_text(aes(x, y, label=labs, group=NULL),data=dat,size=6)
  dat <- data.frame(x = rep(1, len), y = rep(1, len), vars, labs=c("","B",""))
  dat[2, 1:2] <- c(0.75,3.6); p<-p + geom_text(aes(x, y, label=labs, group=NULL),data=dat,size=5)
  p

  # Forest
  dat <- data.frame(x = rep(1, len), y = rep(1, len), vars, labs=c("","","a")) # side dic
  dat[3, 1:2] <- c(1, 2.1); p<-p + geom_text(aes(x, y, label=labs, group=NULL),data=dat)
  dat <- data.frame(x = rep(1, len), y = rep(1, len), vars, labs=c("","","b")) # front dic
  dat[3, 1:2] <- c(2,2.3); p<-p + geom_text(aes(x, y, label=labs, group=NULL),data=dat)
  dat <- data.frame(x = rep(1, len), y = rep(1, len), vars, labs=c("","","b")) # back dic
  dat[3, 1:2] <- c(3, 2.4); p<-p + geom_text(aes(x, y, label=labs, group=NULL),data=dat)
  dat <- data.frame(x = rep(1, len), y = rep(1, len), vars, labs=c("","","c")) # hide dic
  dat[3, 1:2] <- c(4, 2.9); p<-p + geom_text(aes(x, y, label=labs, group=NULL),data=dat)
  dat <- data.frame(x = rep(1, len), y = rep(1, len), vars, labs=c("","","*"))
  dat[3, 1:2] <- c(1,2.2); p<-p + geom_text(aes(x, y, label=labs, group=NULL),data=dat,size=6)
  dat <- data.frame(x = rep(1, len), y = rep(1, len), vars, labs=c("","","*"))
  dat[3, 1:2] <- c(2,2.4); p<-p + geom_text(aes(x, y, label=labs, group=NULL),data=dat,size=6)
  dat <- data.frame(x = rep(1, len), y = rep(1, len), vars, labs=c("","","*"))
  dat[3, 1:2] <- c(3,2.6); p<-p + geom_text(aes(x, y, label=labs, group=NULL),data=dat,size=6)
  dat <- data.frame(x = rep(1, len), y = rep(1, len), vars, labs=c("","","*"))
  dat[3, 1:2] <- c(4,3); p<-p + geom_text(aes(x, y, label=labs, group=NULL),data=dat,size=6)
  dat <- data.frame(x = rep(1, len), y = rep(1, len), vars, labs=c("","","C"))
  dat[3, 1:2] <- c(0.75,3.6); p<-p + geom_text(aes(x, y, label=labs, group=NULL),data=dat,size=5)
  p}
dev.off()

#####
# Figure 2 - Latency x Carnivorous species
#####
df<-na.omit(d1)
levels(df$Predator)
df$Predator<- relevel(df$Predator,ref="ocelot")
df$Predator<- relevel(df$Predator,ref="cougar")
levels(df$Predator)
levels(df$StimulusPosition)
levels(df$StimulusPosition) <- c("Lateral","Anterior","Posterior","Hidden")
levels(df$VisualPhenotype)<-c("Dichromats","Trichromats")
cbPalette <- c("#999999", "#E69F00")
pdf("Fig. 2 - Latency x Carnivorous species.pdf",width=9,height=5)

{p<-ggbarplot(df,x="StimulusPosition", y = "loglat", add=c("mean_se"),facet.by = c("Predator"),
              panel.labs=list(Predator=c("Cougar","Ocelot","Lesser grison")),color = "black", palette = cbPalette,
              fill="VisualPhenotype",size=0.4,ylim=c(0,4), position = position_dodge(0.8)
)
  p<-p+labs(y="Response latency (s, log)",x="Body posture")
  p<-p+theme(legend.title = element_blank())
  p<-p+theme(strip.background =element_rect(fill="white"))
  p<-p+theme(legend.position = c(0.5, 0.93),legend.direction = "horizontal")
  df$Predator<-factor(df$Predator)
  len <- length(levels(df$Predator))
  vars <- data.frame(expand.grid(levels(df$Predator)))
  colnames(vars) <- c("Predator")

  # Cougar
  dat <- data.frame(x = rep(1, len), y = rep(1, len), vars, labs=c("a","","")) # side dic
  dat[1, 1:2] <- c(0.8, 1.4); p<-p + geom_text(aes(x, y, label=labs, group=NULL),data=dat)
  dat <- data.frame(x = rep(1, len), y = rep(1, len), vars, labs=c("a","","")) # front dic
  dat[1, 1:2] <- c(1.8, 1.4); p<-p + geom_text(aes(x, y, label=labs, group=NULL),data=dat)
  dat <- data.frame(x = rep(1, len), y = rep(1, len), vars, labs=c("b","","")) # back dic
  dat[1, 1:2] <- c(2.8, 1.8); p<-p + geom_text(aes(x, y, label=labs, group=NULL),data=dat)
  dat <- data.frame(x = rep(1, len), y = rep(1, len), vars, labs=c("c","","")) # hide dic
  dat[1, 1:2] <- c(3.8, 2.3); p<-p + geom_text(aes(x, y, label=labs, group=NULL),data=dat)
  dat <- data.frame(x = rep(1, len), y = rep(1, len), vars, labs=c("A","","")) # side dic
  dat[1, 1:2] <- c(1.2, 0.9); p<-p + geom_text(aes(x, y, label=labs, group=NULL),data=dat)
  dat <- data.frame(x = rep(1, len), y = rep(1, len), vars, labs=c("AB","","")) # front dic
  dat[1, 1:2] <- c(2.2, 1); p<-p + geom_text(aes(x, y, label=labs, group=NULL),data=dat)
  dat <- data.frame(x = rep(1, len), y = rep(1, len), vars, labs=c("B","","")) # back dic
  dat[1, 1:2] <- c(3.2, 1.3); p<-p + geom_text(aes(x, y, label=labs, group=NULL),data=dat)
  dat <- data.frame(x = rep(1, len), y = rep(1, len), vars, labs=c("C","","")) # hide dic
  dat[1, 1:2] <- c(4.2, 1.7); p<-p + geom_text(aes(x, y, label=labs, group=NULL),data=dat)
  dat <- data.frame(x = rep(1, len), y = rep(1, len), vars, labs=c("*","",""))
  dat[1, 1:2] <- c(1,1.5); p<-p + geom_text(aes(x, y, label=labs, group=NULL),data=dat,size=6)
  dat <- data.frame(x = rep(1, len), y = rep(1, len), vars, labs=c("*","",""))
  dat[1, 1:2] <- c(2,1.5); p<-p + geom_text(aes(x, y, label=labs, group=NULL),data=dat,size=6)
  dat <- data.frame(x = rep(1, len), y = rep(1, len), vars, labs=c("*","",""))
  dat[1, 1:2] <- c(3,1.8); p<-p + geom_text(aes(x, y, label=labs, group=NULL),data=dat,size=6)
  dat <- data.frame(x = rep(1, len), y = rep(1, len), vars, labs=c("*","",""))
  dat[1, 1:2] <- c(4,2.3); p<-p + geom_text(aes(x, y, label=labs, group=NULL),data=dat,size=6)
  dat <- data.frame(x = rep(1, len), y = rep(1, len), vars, labs=c("A","",""))
  dat[1, 1:2] <- c(0.75,3.6); p<-p + geom_text(aes(x, y, label=labs, group=NULL),data=dat,size=5)
  p

  # Ocelot
  dat <- data.frame(x = rep(1, len), y = rep(1, len), vars, labs=c("","a","")) # side dic
  dat[2, 1:2] <- c(1, 2); p<-p + geom_text(aes(x, y, label=labs, group=NULL),data=dat)
  dat <- data.frame(x = rep(1, len), y = rep(1, len), vars, labs=c("","a","")) # front dic
  dat[2, 1:2] <- c(2,2.1); p<-p + geom_text(aes(x, y, label=labs, group=NULL),data=dat)
  dat <- data.frame(x = rep(1, len), y = rep(1, len), vars, labs=c("","a","")) # back dic
  dat[2, 1:2] <- c(3, 2.1); p<-p + geom_text(aes(x, y, label=labs, group=NULL),data=dat)
  dat <- data.frame(x = rep(1, len), y = rep(1, len), vars, labs=c("","b","")) # hide dic
  dat[2, 1:2] <- c(4, 2.9); p<-p + geom_text(aes(x, y, label=labs, group=NULL),data=dat)
  dat <- data.frame(x = rep(1, len), y = rep(1, len), vars, labs=c("","",""))
  dat[2, 1:2] <- c(1,1.8); p<-p + geom_text(aes(x, y, label=labs, group=NULL),data=dat,size=6)
  dat <- data.frame(x = rep(1, len), y = rep(1, len), vars, labs=c("","*",""))
  dat[2, 1:2] <- c(2,2.3); p<-p + geom_text(aes(x, y, label=labs, group=NULL),data=dat,size=6)
  dat <- data.frame(x = rep(1, len), y = rep(1, len), vars, labs=c("","*",""))
  dat[2, 1:2] <- c(3,2.2); p<-p + geom_text(aes(x, y, label=labs, group=NULL),data=dat,size=6)
  dat <- data.frame(x = rep(1, len), y = rep(1, len), vars, labs=c("","*",""))
  dat[2, 1:2] <- c(4,3.2); p<-p + geom_text(aes(x, y, label=labs, group=NULL),data=dat,size=6)
  dat <- data.frame(x = rep(1, len), y = rep(1, len), vars, labs=c("","B",""))
  dat[2, 1:2] <- c(0.75,3.6); p<-p + geom_text(aes(x, y, label=labs, group=NULL),data=dat,size=5)
  p

  # Lesser grison
  dat <- data.frame(x = rep(1, len), y = rep(1, len), vars, labs=c("","","a")) # side dic
  dat[3, 1:2] <- c(1, 1.9); p<-p + geom_text(aes(x, y, label=labs, group=NULL),data=dat)
  dat <- data.frame(x = rep(1, len), y = rep(1, len), vars, labs=c("","","a")) # front dic
  dat[3, 1:2] <- c(2,1.9); p<-p + geom_text(aes(x, y, label=labs, group=NULL),data=dat)
  dat <- data.frame(x = rep(1, len), y = rep(1, len), vars, labs=c("","","b")) # back dic
  dat[3, 1:2] <- c(3, 2.9); p<-p + geom_text(aes(x, y, label=labs, group=NULL),data=dat)
  dat <- data.frame(x = rep(1, len), y = rep(1, len), vars, labs=c("","","c")) # hide dic
  dat[3, 1:2] <- c(4, 3.1); p<-p + geom_text(aes(x, y, label=labs, group=NULL),data=dat)
  dat <- data.frame(x = rep(1, len), y = rep(1, len), vars, labs=c("","",""))
  dat[3, 1:2] <- c(1,1.6); p<-p + geom_text(aes(x, y, label=labs, group=NULL),data=dat,size=6)
  dat <- data.frame(x = rep(1, len), y = rep(1, len), vars, labs=c("","",""))
  dat[3, 1:2] <- c(2,1.4); p<-p + geom_text(aes(x, y, label=labs, group=NULL),data=dat,size=6)
  dat <- data.frame(x = rep(1, len), y = rep(1, len), vars, labs=c("","",""))
  dat[3, 1:2] <- c(3,2); p<-p + geom_text(aes(x, y, label=labs, group=NULL),data=dat,size=6)
  dat <- data.frame(x = rep(1, len), y = rep(1, len), vars, labs=c("","",""))
  dat[3, 1:2] <- c(4,2.6); p<-p + geom_text(aes(x, y, label=labs, group=NULL),data=dat,size=6)
  dat <- data.frame(x = rep(1, len), y = rep(1, len), vars, labs=c("","","C"))
  dat[3, 1:2] <- c(0.75,3.6); p<-p + geom_text(aes(x, y, label=labs, group=NULL),data=dat,size=5)
  p}
dev.off()

#####
# Figure 3 - Latency x Stimulus dimension
#####
df<-na.omit(d1)
levels(df$Size)
df$Size<- relevel(df$Size,ref="large")
levels(df$Size)
levels(df$StimulusPosition)
levels(df$StimulusPosition) <- c("Lateral","Anterior","Posterior","Hidden")
levels(df$VisualPhenotype)<-c("Dichromats","Trichromats")
cbPalette <- c("#999999", "#E69F00")
pdf("Fig. 3 - Latency x Stimulus dimension.pdf",width=9,height=5)

{p<-ggbarplot(df,x="StimulusPosition", y = "loglat", add=c("mean_se"),facet.by = c("Size"),
              panel.labs=list(Size=c("Large","Small")),color = "black", palette = cbPalette,
              fill="VisualPhenotype",size=0.4,ylim=c(0,4), position = position_dodge(0.8)
)
  p<-p+labs(y="Response latency (s, log)",x="Body posture")
  p<-p+theme(legend.title = element_blank())
  p<-p+theme(strip.background =element_rect(fill="white"))
  p<-p+theme(legend.position = c(0.25, 0.93),legend.direction = "horizontal")
  df$Size<-factor(df$Size)
  len <- length(levels(df$Size))
  vars <- data.frame(expand.grid(levels(df$Size)))
  colnames(vars) <- c("Size")

  # Large
  dat <- data.frame(x = rep(1, len), y = rep(1, len), vars, labs=c("a","")) # side dic
  dat[1, 1:2] <- c(1, 1.5);  p<-p + geom_text(aes(x, y, label=labs, group=NULL),data=dat)
  dat <- data.frame(x = rep(1, len), y = rep(1, len), vars, labs=c("b","")) # front dic
  dat[1, 1:2] <- c(2, 1.7);  p<-p + geom_text(aes(x, y, label=labs, group=NULL),data=dat)
  dat <- data.frame(x = rep(1, len), y = rep(1, len), vars, labs=c("c","")) # back dic
  dat[1, 1:2] <- c(3, 2);  p<-p + geom_text(aes(x, y, label=labs, group=NULL),data=dat)
  dat <- data.frame(x = rep(1, len), y = rep(1, len), vars, labs=c("d","")) # hide dic
  dat[1, 1:2] <- c(4, 2.8);  p<-p + geom_text(aes(x, y, label=labs, group=NULL),data=dat)
  dat <- data.frame(x = rep(1, len), y = rep(1, len), vars, labs=c("*",""))
  dat[1, 1:2] <- c(1,1.7);  p<-p + geom_text(aes(x, y, label=labs, group=NULL),data=dat,size=6)
  dat <- data.frame(x = rep(1, len), y = rep(1, len), vars, labs=c("*",""))
  dat[1, 1:2] <- c(2,1.9);  p<-p + geom_text(aes(x, y, label=labs, group=NULL),data=dat,size=6)
  dat <- data.frame(x = rep(1, len), y = rep(1, len), vars, labs=c("*",""))
  dat[1, 1:2] <- c(3,2.1);  p<-p + geom_text(aes(x, y, label=labs, group=NULL),data=dat,size=6)
  dat <- data.frame(x = rep(1, len), y = rep(1, len), vars, labs=c("*",""))
  dat[1, 1:2] <- c(4,3);  p<-p + geom_text(aes(x, y, label=labs, group=NULL),data=dat,size=6)
  dat <- data.frame(x = rep(1, len), y = rep(1, len), vars, labs=c("A",""))
  dat[1, 1:2] <- c(0.75,4);  p<-p + geom_text(aes(x, y, label=labs, group=NULL),data=dat,size=5)
  p

  # Small
  dat <- data.frame(x = rep(1, len), y = rep(1, len), vars, labs=c("","a")) # side dic
  dat[2, 1:2] <- c(1,1.9);  p<-p + geom_text(aes(x, y, label=labs, group=NULL),data=dat)
  dat <- data.frame(x = rep(1, len), y = rep(1, len), vars, labs=c("","a")) # front dic
  dat[2, 1:2] <- c(2,1.8);  p<-p + geom_text(aes(x, y, label=labs, group=NULL),data=dat)
  dat <- data.frame(x = rep(1, len), y = rep(1, len), vars, labs=c("","b")) # back dic
  dat[2, 1:2] <- c(3,2.4);  p<-p + geom_text(aes(x, y, label=labs, group=NULL),data=dat)
  dat <- data.frame(x = rep(1, len), y = rep(1, len), vars, labs=c("","c")) # hide dic
  dat[2, 1:2] <- c(4,2.8);  p<-p + geom_text(aes(x, y, label=labs, group=NULL),data=dat)
  dat <- data.frame(x = rep(1, len), y = rep(1, len), vars, labs=c("","*"))
  dat[2, 1:2] <- c(1,2.2);  p<-p + geom_text(aes(x, y, label=labs, group=NULL),data=dat,size=6)
  dat <- data.frame(x = rep(1, len), y = rep(1, len), vars, labs=c("","*"))
  dat[2, 1:2] <- c(2,1.9);  p<-p + geom_text(aes(x, y, label=labs, group=NULL),data=dat,size=6)
  dat <- data.frame(x = rep(1, len), y = rep(1, len), vars, labs=c("","*"))
  dat[2, 1:2] <- c(3,2.6);  p<-p + geom_text(aes(x, y, label=labs, group=NULL),data=dat,size=6)
  dat <- data.frame(x = rep(1, len), y = rep(1, len), vars, labs=c("","*"))
  dat[2, 1:2] <- c(4,3);  p<-p + geom_text(aes(x, y, label=labs, group=NULL),data=dat,size=6)
  dat <- data.frame(x = rep(1, len), y = rep(1, len), vars, labs=c("","B"))
  dat[2, 1:2] <- c(0.75,4);  p<-p + geom_text(aes(x, y, label=labs, group=NULL),data=dat,size=5)
  p}
dev.off()

#####
# Figure 4 - Detection (3s) x Background scenario
#####
levels(d1$StimulusPosition)
levels(d1$StimulusPosition) <- c("Lateral","Anterior","Posterior","Hidden")
levels(df$VisualPhenotype)<-c("Dichromats","Trichromats")
pdf("Fig. 4 - Detection (3s) x Background scenario.pdf",width=10,height=5,onefile=F)
{d2<-subset(df,Background=="grassland")
  d2<-na.omit(d2)
  levels(d2$Background)
  m2<-glmmTMB(Detection~StimulusPosition+VisualPhenotype+Predator+Size+(1|Subject),family="binomial"(link=probit),data=d2)
  p1<-plot_model(m2,type="emm",terms=c("StimulusPosition","VisualPhenotype"),ci.lvl=0.95,grid=F,legend="VisualPhenotype",
                 x.cat = TRUE,title="",colors=c("#999999", "#E69F00"),dodge=0.7,dot.size=4)
  p1<-p1+labs(x="Body posture",y="Detection accuracy (3s)")
  p1<-p1+theme_classic()
  p1<-p1+theme(axis.text.x=element_text(hjust=0.5,size=9),axis.title = element_text(color = "black"))
  p1<-p1+annotate("text", x = 1, y = 0.73, label = "a") # side tri
  p1<-p1+annotate("text", x = 2, y = 0.85, label = "b") # front tri
  p1<-p1+annotate("text", x = 3, y = 0.57, label = "c") # back tri
  p1<-p1+annotate("text", x = 4, y = 0.32, label = "d") # hide tri
  p1<-p1+annotate("text", x = 1, y = 0.76, label = "*",size=6)
  p1<-p1+annotate("text", x = 2, y = 0.88, label = "*",size=6)
  p1<-p1+annotate("text", x = 3, y = 0.60, label = "*",size=6)
  p1<-p1+annotate("text", x = 4, y = 0.35, label = "*",size=6)
  p1<-p1+annotate("text", x = 0.75, y = 1, label = "A",size=5)
  p1<-p1+set_theme(base=theme_classic(),legend.pos = "top",legend.item.backcol = "white",legend.title.face = "plain",
                   axis.textcolor = "black",geom.outline.color = "black",geom.label.color="black")
  p1<-p1+theme(axis.title = element_text(color = "black"))
  p1<-p1+theme(legend.position = c(0.2, 1),legend.direction = "horizontal")
  p1<-p1+theme(panel.background = element_rect(colour = "black", size=1))
  p1<-p1+theme(legend.title = element_blank())
  p1<-p1+ylim(0,1.05)
  p1<-p1+ggtitle("Grassland")+theme(plot.title = element_text(hjust = 0.5))}

{d2<-subset(df,Background=="savanna")
  d2<-na.omit(d2)
  levels(d2$Background)
  m1<-glmmTMB(Detection~StimulusPosition*VisualPhenotype+Predator+Size+(1|Subject),family="binomial"(link=logit),data=d2)
  p2<-plot_model(m1,type="emm",terms=c("StimulusPosition","VisualPhenotype"),ci.lvl=0.95,grid=F,legend="VisualPhenotype",
                 x.cat = TRUE,title="",colors=c("#999999", "#E69F00"),dodge=0.7,dot.size=4)
  p2<-p2+labs(x="Body posture",y="Detection accuracy (3s)")
  p2<-p2+theme_classic()
  p2<-p2+theme(axis.text.x=element_text(hjust=0.5,size=9),axis.title = element_text(color = "black"))
  p2<-p2+annotate("text", x = 1, y = 0.63, label = "a") # side tri
  p2<-p2+annotate("text", x = 2, y = 0.6, label = "a") # front tri
  p2<-p2+annotate("text", x = 3, y = 0.4, label = "b") # back tri
  p2<-p2+annotate("text", x = 4, y = 0.15, label = "c") # hide tri
  p2<-p2+annotate("text", x = 1, y = 0.66, label = "*",size=6)
  p2<-p2+annotate("text", x = 2, y = 0.63, label = "*",size=6)
  p2<-p2+annotate("text", x = 3, y = 0.43, label = "*",size=6)
  p2<-p2+annotate("text", x = 4, y = 0.18, label = "*",size=6)
  p2<-p2+annotate("text", x = 0.75, y = 1, label = "B",size=5)
  p2<-p2+set_theme(base=theme_classic(),legend.pos = "top",legend.item.backcol = "white",legend.title.face = "plain",
                   axis.textcolor = "black",geom.outline.color = "black",geom.label.color="black")
  p2<-p2+theme(axis.title = element_text(color = "black"))
  p2<-p2+theme(legend.position = c(0.2, 1),legend.direction = "horizontal")
  p2<-p2+theme(panel.background = element_rect(colour = "black", size=1))
  p2<-p2+theme(legend.title = element_blank())
  p2<-p2+ylim(0,1.05)
  p2<-p2+ggtitle("Savanna")+theme(plot.title = element_text(hjust = 0.5))}

{d2<-subset(df,Background=="forest")
  d2<-na.omit(d2)
  levels(d2$Background)
  m3<-glmmTMB(Detection~StimulusPosition+VisualPhenotype+Predator+Size+(1|Subject),family="binomial"(link=cloglog),data=d2)
  p3<-plot_model(m3,type="emm",terms=c("StimulusPosition","VisualPhenotype"),ci.lvl=0.95,grid=F,legend="VisualPhenotype",
                 x.cat = TRUE,title="",colors=c("#999999", "#E69F00"),dodge=0.7,dot.size=4)
  p3<-p3+labs(x="Body posture",y="Detection accuracy (3s)")
  p3<-p3+theme_classic()
  p3<-p3+theme(axis.text.x=element_text(hjust=0.5,size=9),axis.title = element_text(color = "black"))
  p3<-p3+annotate("text", x = 1, y = 0.33, label = "a") # side tri
  p3<-p3+annotate("text", x = 2, y = 0.29, label = "ab") # front tri
  p3<-p3+annotate("text", x = 3, y = 0.25, label = "b") # back tri
  p3<-p3+annotate("text", x = 4, y = 0.2, label = "c") # hide tri
  p3<-p3+annotate("text", x = 1, y = 0.37, label = "*",size=6)
  p3<-p3+annotate("text", x = 2, y = 0.32, label = "*",size=6)
  p3<-p3+annotate("text", x = 3, y = 0.28, label = "*",size=6)
  p3<-p3+annotate("text", x = 4, y = 0.23, label = "*",size=6)
  p3<-p3+annotate("text", x = 0.75, y = 1, label = "C",size=5)
  p3<-p3+set_theme(base=theme_classic(),legend.pos = "top",legend.item.backcol = "white",legend.title.face = "plain",
                   axis.textcolor = "black",geom.outline.color = "black",geom.label.color="black")
  p3<-p3+theme(axis.title = element_text(color = "black"))
  p3<-p3+theme(legend.position = c(0.2, 1),legend.direction = "horizontal")
  p3<-p3+theme(panel.background = element_rect(colour = "black", size=1))
  p3<-p3+theme(legend.title = element_blank())
  p3<-p3+ylim(0,1.05)
  p3<-p3+ggtitle("Forest")+theme(plot.title = element_text(hjust = 0.5))}

p4<-ggarrange(p1,p2,p3,ncol=3,nrow=1,label.x=0.95,label.y=0.9,
              common.legend = TRUE, legend = "top")
p4
dev.off()

#####
# Figure 5 - Detection (3s) x Carnivorous species
#####
pdf("Fig. 5 - Detection (3s) x Carnivorous species.pdf",width=10,height=5,onefile=F)
levels(d1$StimulusPosition)
levels(d1$StimulusPosition) <- c("Lateral","Anterior","Posterior","Hidden")
levels(d1$VisualPhenotype)<-c("Dichromats","Trichromats")
{d2<-subset(d1,Predator=="lesser grison")
  d2<-na.omit(d2)
  levels(d2$Predator)
  m3<-glmmTMB(Detection~StimulusPosition+VisualPhenotype+Background+Size+(1|Subject),family="binomial"(link=cloglog),data=d2)
  p1<-plot_model(m3,type="emm",terms=c("StimulusPosition","VisualPhenotype"),ci.lvl=0.95,grid=F,legend="VisualPhenotype",
                 x.cat = TRUE,title="",colors=c("#999999", "#E69F00"),dodge=0.7,dot.size=4)
  p1<-p1+labs(x="Body posture",y="Detection accuracy (3s)")
  p1<-p1+theme_classic()
  p1<-p1+theme(axis.text.x=element_text(hjust=0.5,size=9),axis.title = element_text(color = "black"))
  p1<-p1+annotate("text", x = 1, y = 0.40, label = "a") # side tri
  p1<-p1+annotate("text", x = 2, y = 0.35, label = "a") # front tri
  p1<-p1+annotate("text", x = 3, y = 0.15, label = "b") # back tri
  p1<-p1+annotate("text", x = 4, y = 0.10, label = "c") # hide tri
  p1<-p1+annotate("text", x = 0.75, y = 1, label = "C",size=5)
  p1<-p1+set_theme(base=theme_classic(),legend.pos = "top",legend.item.backcol = "white",legend.title.face = "plain",
                   axis.textcolor = "black",geom.outline.color = "black",geom.label.color="black")
  p1<-p1+theme(axis.title = element_text(color = "black"))
  p1<-p1+theme(legend.position = c(0.15, 0.5),legend.direction = "horizontal")
  p1<-p1+theme(panel.background = element_rect(colour = "black", size=1))
  p1<-p1+theme(legend.title = element_blank())
  p1<-p1+ylim(0,1.05)
  p1<-p1+ggtitle("Lesser grison")+theme(plot.title = element_text(hjust = 0.5))}

{d2<-subset(d1,Predator=="ocelot")
  d2<-na.omit(d2)
  levels(d2$Predator)
  m3<-glmmTMB(Detection~StimulusPosition+VisualPhenotype+Background+Size+(1|Subject),family="binomial"(link=cloglog),data=d2)
  p2<-plot_model(m3,type="emm",terms=c("StimulusPosition","VisualPhenotype"),ci.lvl=0.95,grid=F,legend="VisualPhenotype",
                 x.cat = TRUE,title="",colors=c("#999999", "#E69F00"),dodge=0.7,dot.size=4)
  p2<-p2+labs(x="Body posture",y="Detection accuracy (3s)")
  p2<-p2+theme_classic()
  p2<-p2+theme(axis.text.x=element_text(hjust=0.5,size=9),axis.title = element_text(color = "black"))
  p2<-p2+annotate("text", x = 1, y = 0.35, label = "ab") # side tri
  p2<-p2+annotate("text", x = 2, y = 0.45, label = "a") # front tri
  p2<-p2+annotate("text", x = 3, y = 0.3, label = "b") # back tri
  p2<-p2+annotate("text", x = 4, y = 0.15, label = "c") # hide tri
  p2<-p2+annotate("text", x = 1, y = 0.38, label = "*",size=6)
  p2<-p2+annotate("text", x = 2, y = 0.48, label = "*",size=6)
  p2<-p2+annotate("text", x = 3, y = 0.33, label = "*",size=6)
  p2<-p2+annotate("text", x = 4, y = 0.18, label = "*",size=6)
  p2<-p2+annotate("text", x = 0.75, y = 1, label = "B",size=5)
  p2<-p2+set_theme(base=theme_classic(),legend.pos = "top",legend.item.backcol = "white",legend.title.face = "plain",
                   axis.textcolor = "black",geom.outline.color = "black",geom.label.color="black")
  p2<-p2+theme(axis.title = element_text(color = "black"))
  p2<-p2+theme(legend.position = c(0.15, 0.5),legend.direction = "horizontal")
  p2<-p2+theme(panel.background = element_rect(colour = "black", size=1))
  p2<-p2+theme(legend.title = element_blank())
  p2<-p2+ylim(0,1.05)
  p2<-p2+ggtitle("Ocelot")+theme(plot.title = element_text(hjust = 0.5))}

{d2<-subset(d1,Predator=="cougar")
  d2<-na.omit(d2)
  levels(d2$Predator)
  m1<-glmmTMB(Detection~StimulusPosition+VisualPhenotype+Background+Size+(1|Subject),family="binomial"(link=logit),data=d2)
  p3<-plot_model(m1,type="emm",terms=c("StimulusPosition","VisualPhenotype"),ci.lvl=0.95,grid=F,legend="VisualPhenotype",
                 x.cat = TRUE,title="",colors=c("#999999", "#E69F00"),dodge=0.7,dot.size=4)
  p3<-p3+labs(x="Body posture",y="Detection accuracy (3s)")
  p3<-p3+theme_classic()
  p3<-p3+theme(axis.text.x=element_text(hjust=0.5,size=9),axis.title = element_text(color = "black"))
  p3<-p3+annotate("text", x = 1, y = 0.9, label = "a") # side tri
  p3<-p3+annotate("text", x = 2, y = 0.9, label = "a") # front tri
  p3<-p3+annotate("text", x = 3, y = 0.8, label = "b") # back tri
  p3<-p3+annotate("text", x = 4, y = 0.45, label = "c") # hide tri
  p3<-p3+annotate("text", x = 1, y = 0.93, label = "*",size=6)
  p3<-p3+annotate("text", x = 2, y = 0.93, label = "*",size=6)
  p3<-p3+annotate("text", x = 3, y = 0.83, label = "*",size=6)
  p3<-p3+annotate("text", x = 4, y = 0.48, label = "*",size=6)
  p3<-p3+annotate("text", x = 0.75, y = 1, label = "A",size=5)
  p3<-p3+set_theme(base=theme_classic(),legend.pos = "top",legend.item.backcol = "white",legend.title.face = "plain",
                   axis.textcolor = "black",geom.outline.color = "black",geom.label.color="black")
  p3<-p3+theme(axis.title = element_text(color = "black"))
  p3<-p3+theme(legend.position = c(0.15, 0.5),legend.direction = "horizontal")
  p3<-p3+theme(panel.background = element_rect(colour = "black", size=1))
  p3<-p3+theme(legend.title = element_blank())
  p3<-p3+ylim(0,1.05)
  p3<-p3+ggtitle("Cougar")+theme(plot.title = element_text(hjust = 0.5))}

p4<-ggarrange(p3,p2,p1,ncol=3,nrow=1,label.x=0.95,label.y=0.9,
              common.legend = TRUE, legend = "top")
p4
dev.off()

#####
# Figure 6 - Detection (3s) x Stimulus dimension
#####
levels(d1$StimulusPosition)
levels(d1$StimulusPosition) <- c("Lateral","Anterior","Posterior","Hidden")
levels(d1$VisualPhenotype)<-c("Dichromats","Trichromats")
pdf("Fig. 6 - Detection (3s) x Stimulus dimension.pdf",width=8,height=5,onefile=F)
{d2<-subset(d1,Size=="large")
  d2<-na.omit(d2)
  levels(d2$Size)
  m3<-glmmTMB(Detection~StimulusPosition*VisualPhenotype+Background+Predator+(1|Subject),family="binomial"(link=cloglog),data=d2)
  p1<-plot_model(m3,type="emm",terms=c("StimulusPosition","VisualPhenotype"),ci.lvl=0.95,grid=F,legend="VisualPhenotype",
                 x.cat = TRUE,title="",colors=c("#999999", "#E69F00"),dodge=0.7,dot.size=4)
  p1<-p1+labs(x="Body posture",y="Detection accuracy (3s)")
  p1<-p1+theme_classic()
  p1<-p1+theme(axis.text.x=element_text(hjust=0.5,size=9),axis.title = element_text(color = "black"))
  p1<-p1+annotate("text", x = 0.825, y = 0.55, label = "a") # side dic
  p1<-p1+annotate("text", x = 1.825, y = 0.5, label = "a") # front dic
  p1<-p1+annotate("text", x = 2.825, y = 0.3, label = "b") # back dic
  p1<-p1+annotate("text", x = 3.825, y = 0.15, label = "c") # hide dic
  p1<-p1+annotate("text", x = 1.175, y = 0.65, label = "A") # side tri
  p1<-p1+annotate("text", x = 2.175, y = 0.55, label = "AB") # front tri
  p1<-p1+annotate("text", x = 3.175, y = 0.5, label = "B") # back tri
  p1<-p1+annotate("text", x = 4.175, y = 0.22, label = "C") # hide tri
  p1<-p1+annotate("text", x = 1, y = 0.37, label = "",size=6)
  p1<-p1+annotate("text", x = 2, y = 0.47, label = "",size=6)
  p1<-p1+annotate("text", x = 3, y = 0.5, label = "*",size=6)
  p1<-p1+annotate("text", x = 4, y = 0.25, label = "*",size=6)
  p1<-p1+annotate("text", x = 0.75, y = 1, label = "A",size=5)
  p1<-p1+set_theme(base=theme_classic(),legend.pos = "top",legend.item.backcol = "white",legend.title.face = "plain",
                   axis.textcolor = "black",geom.outline.color = "black",geom.label.color="black")
  p1<-p1+theme(axis.title = element_text(color = "black"))
  p1<-p1+theme(legend.position = c(0.15, 0.8),legend.direction = "horizontal")
  p1<-p1+theme(panel.background = element_rect(colour = "black", size=1))
  p1<-p1+theme(legend.title = element_blank())
  p1<-p1+ylim(0,1.05)
  p1<-p1+ggtitle("Large")+theme(plot.title = element_text(hjust = 0.5))}

{d2<-subset(d1,Size=="small")
  d2<-na.omit(d2)
  levels(d2$Size)
  m1<-glmmTMB(Detection~StimulusPosition+VisualPhenotype+Background+Predator+(1|Subject),family="binomial"(link=logit),data=d2)
  p2<-plot_model(m1,type="emm",terms=c("StimulusPosition","VisualPhenotype"),ci.lvl=0.95,grid=F,legend="VisualPhenotype",
                 x.cat = TRUE,title="",colors=c("#999999", "#E69F00"),dodge=0.7,dot.size=4)
  p2<-p2+labs(x="Body posture",y="Detection accuracy (3s)")
  p2<-p2+theme_classic()
  p2<-p2+theme(axis.text.x=element_text(hjust=0.5,size=9),axis.title = element_text(color = "black"))
  p2<-p2+annotate("text", x = 1, y = 0.5, label = "a") # side tri
  p2<-p2+annotate("text", x = 2, y = 0.65, label = "b") # front tri
  p2<-p2+annotate("text", x = 3, y = 0.3, label = "c") # back tri
  p2<-p2+annotate("text", x = 4, y = 0.2, label = "d") # hide tri
  p2<-p2+annotate("text", x = 1, y = 0.53, label = "*",size=6)
  p2<-p2+annotate("text", x = 2, y = 0.68, label = "*",size=6)
  p2<-p2+annotate("text", x = 3, y = 0.33, label = "*",size=6)
  p2<-p2+annotate("text", x = 4, y = 0.23, label = "*",size=6)
  p2<-p2+annotate("text", x = 0.75, y = 1, label = "B",size=5)
  p2<-p2+set_theme(base=theme_classic(),legend.pos = "top",legend.item.backcol = "white",legend.title.face = "plain",
                   axis.textcolor = "black",geom.outline.color = "black",geom.label.color="black")
  p2<-p2+theme(axis.title = element_text(color = "black"))
  p2<-p2+theme(legend.position = c(0.15, 0.9),legend.direction = "horizontal")
  p2<-p2+theme(panel.background = element_rect(colour = "black", size=1))
  p2<-p2+theme(legend.title = element_blank())
  p2<-p2+ylim(0,1.05)
  p2<-p2+ggtitle("Small")+theme(plot.title = element_text(hjust = 0.5))}

p3<-ggarrange(p1,p2,ncol=2,nrow=1,label.x=0.95,label.y=0.9,
              common.legend = TRUE, legend = "top")
p3
dev.off()

#####
# Figure 7 - Latency + Detection (3s) x Carnivore species + Visual phenotype
#####

# in pdfsam basic delete 1st black page generated by the pdf:

df<-na.omit(d1)
levels(df$StimulusPosition)
levels(df$StimulusPosition) <- c("Lateral","Anterior","Posterior","Hidden")
levels(df$VisualPhenotype)<-c("Dichromats","Trichromats")
df<-subset(df,StimulusPosition=="Anterior")
df$Predator<- relevel(df$Predator,ref="Ocelot")
df$Predator<- relevel(df$Predator,ref="Cougar")
str(df)
cbPalette <- c("#999999", "#E69F00")

pdf("Fig. 7 - Latency and detection x carnivore and phenotype.pdf",width=8,height=5)
{p<-ggbarplot(df,x="Predator", y = "loglat", add=c("mean_se"),color = "black", palette = cbPalette,
              fill="VisualPhenotype",size=0.4,ylim=c(0,4), position = position_dodge(0.8)
)
  p<-p+labs(y="Response latency (s, log)",x="Carnivore model")
  p<-p+theme(legend.title = element_blank())
  p<-p+theme(strip.background =element_rect(fill="black"))
  p<-p+theme(legend.position = c(0.5, 0.93),legend.direction = "horizontal")
  p<-p+theme(panel.background = element_rect(colour = "black", size=1))
  p<-p+annotate("text",x=1,y=1.4,label="a")
  p<-p+annotate("text",x=2,y=2.1,label="b")
  p<-p+annotate("text",x=3,y=2.1,label="b")
  p<-p+annotate("text",x=1,y=1.8,size=6,label="*")
  p<-p+annotate("text",x=2,y=2.5,size=6,label="*")
  p<-p+annotate("text", x = 0.65, y = 3.9, label = "A",size=5)}

{d2<-df
  levels(d2$Size)
  m3<-glmmTMB(Detection~Predator*VisualPhenotype+Background+Size+(1|Subject),family="binomial"(link=probit),data=d2)
  p1<-plot_model(m3,type="emm",terms=c("Predator","VisualPhenotype"),ci.lvl=0.95,grid=F,legend="VisualPhenotype",
                 x.cat = TRUE,title="",colors=c("#999999", "#E69F00"),dodge=0.7,dot.size=4)
  p1<-p1+labs(x="Carnivore model",y="Detection accuracy (3s)")
  p1<-p1+theme_classic()
  p1<-p1+theme(axis.text.x=element_text(hjust=0.5,size=9),axis.title = element_text(color = "black"))
  p1<-p1+annotate("text", x = 1, y = 0.9, label = "a")
  p1<-p1+annotate("text", x = 2, y = 0.48, label = "b")
  p1<-p1+annotate("text", x = 3, y = 0.46, label = "b")
  p1<-p1+annotate("text", x = 1, y = 0.94, label = "*",size=6)
  p1<-p1+annotate("text", x = 2, y = 0.52, label = "",size=6)
  p1<-p1+annotate("text", x = 3, y = 0.5, label = "",size=6)
  p1<-p1+annotate("text", x = 0.75, y = 1, label = "B",size=5)
  p1<-p1+set_theme(base=theme_classic(),legend.pos = "top",legend.item.backcol = "white",legend.title.face = "plain",
                   axis.textcolor = "black",geom.outline.color = "black",geom.label.color="black")
  p1<-p1+theme(axis.title = element_text(color = "black"))
  p1<-p1+theme(legend.position = c(0.5,0.93),legend.direction = "horizontal")
  p1<-p1+theme(panel.background = element_rect(colour = "black", size=1))
  p1<-p1+theme(legend.title = element_blank())
  p2<-p1+ylim(0,1.05)}
p3<-ggarrange(p,p1,ncol=2,nrow=1,label.x=0.95,label.y=0.9,align = "h",
              common.legend = TRUE, legend = "top")
p3
dev.off()
```
